# Chlorophyll a degradation in Prokaryotes

**DOI:** 10.64898/2026.03.19.712979

**Authors:** Habibu Aliyu, Hendrik Früh, Gunnar Sturm, Anne-Kristin Kaster

## Abstract

Chlorophyll is one of the most abundant pigments on Earth. Although its degradation is well understood in plants, the role of prokaryotes in this process – despite their vast metabolic capabilities – remains unknown. Recent developments in the field of AI-predicted protein structures have opened new avenues for investigating functional homologies between evolutionary-distant organisms previously inaccessible through traditional sequence- or profile-based methods. Here, we present the first evidence of Chlorophyll *a* (Chl *a*) degradation by prokaryotes, discovered through a novel bioinformatic framework which bridges the gap across the domains of life *via* structural alignments of functionally characterised plant proteins, followed by structure similarity graph-based clustering. Metagenomic sequencing data was assembled and binned, yielding over 70,000 medium- to high-quality genomes in total, furthermore publicly available datasets containing genomes from prokaryotic isolates, metagenome-assembled genomes, as well as single-cell genomes were then mined for prokaryotic homologues of Chl *a* degradation genes. Our analysis revealed over 400 genomes from diverse taxonomic groups and habitats that possess a complete pathway, more than 50% stemming from isolates. Additionally, many other genomes harbour partial pathways, suggesting that Chl *a* degradation capabilities are globally widespread across diverse ecosystems. We then validated our *in silico* findings using the model organism *Shewanella acanthi* and confirmed its Chl *a* degradation capability *via* growth experiments, fluorescence spectroscopy and HPLC analyses. Our findings reveal a previously unrecognised pathway in prokaryotes, highlighting the power of structure-based remote homology detection for uncovering metabolic capabilities and evolutionary relationships.

## Introduction

Chlorophylls (Chls) are essential cofactors in the reaction centres of photosystems I and II, absorbing light energy to facilitate electron transport during photosynthesis [1]. They are the most abundant pigments on earth with an estimated annual global production of approximately 1.15 billion tonnes [2], making their formation and degradation one of the most crucial biochemical process on Earth [3]. Chls are a family of cyclic tetrapyrroles with a characteristic isocyclic five-membered ring, consisting of more than 100 different structures, which partake in light capture in photosynthetic organisms [4]. The chemical and physical properties of tetrapyrroles, such as accepting different redox states or absorbing light at specific wavelengths, are governed by the number and location of their conjugated double bonds, side chains and the coordination of different metal ions (Supplementary Fig. S1). Among these diverse pigments, chlorophyll (Chl) *a* is the primary light harvesting and most abundant Chl type in the majority of all oxygenic photosynthetic organisms [5].

### Chlorophyll *a* degradation in plants

Chl *a* degradation is best studied in plants, especially in the angiosperm *Arabidopsis thaliana* during *s*everal plant life-cycle stages. The studied processes include fruit ripening, the steady state cycling due to photosystem damage [6, 7] and the subsequent repair of photosystem II [8]. The PAO/phyllobilin pathway, active during leaf senescence, is currently the best studied process and therefore provides the most comprehensive anabolic scheme for the molecule (Fig. 1) [9–11]. It is fully localised in the chloroplast and typically starts at the dechelation of the magnesium ion from the macrocycle, mediated by Stay-Green1/2 (SGR1/SGR2) [12], yielding pheophytin (Pheo) *a.* Afterwards the phytol tail is hydrolysed by Pheophytinase (PPH) [13] producing pheophorbide (Pheide) *a*. The order of the first two reactions is reversed in young leafs as well as during steady-state cycling due to photosystem damage. Here the phytol tail is hydrolysed first by Chlorophyllase1/2 (CLH1/CLH2) in young leafs [14–16] or by Chlorophyll Dephytylase1 (CLD1) [7] during steady-state cycling, producing chlorophyllide (Chlide) *a* as intermediate. The reaction is followed by the magnesium dechelation through Stay-Green-Like (SGRL), which also yields pheophorbide (Pheide) *a.* The precise function of SGR2 remains disputed, with multiple studies suggesting that the enzyme functions as a negative regulation factor [17, 18], while others claiming it is a stand-by paralog of SGR1 and a positive regulator for the degradation pathway [19]. Since Pheide *a* is a highly active photosensitiser [20] producing cytotoxic reactive oxygen species (ROS) outside the reaction centre of photosystem II if illuminated [21], it needs further detoxification. Pheophorbide *a* oxygenase (PAO), the key enzyme of this degradation pathway, then opens the tetrapyrrole macrocycle of Pheide *a* at the C5 carbon [22, 23] *via* a dioxygenation reaction. The product red chlorophyll catabolite (RCC) is also phototoxic [24] and therefore immediately processed by Red Chlorophyll Catabolite Reductase (RCCR) [25, 26]. Through the reduction of the double bond between C20 and C1 (Supplementary Fig. S1), one of two primary fluorescent chlorophyll catabolite (pFCC) epimers are produced. PAO and RCCR have been shown to interact for efficient RCC processing *via* substrate channelling [24, 27], with PAO only expressed during leaf senescence, and RCCR constitutively expressed in all growth phases [28]. Interestingly, throughout all discovered pathways, the ones localised in the chloroplast are highly conserved among different plants, with only the stereospecificity of RCCR varying across species [24, 29]. As soon as the pFCCs leave the chloroplast, all subsequent modifications are specific to the plant species, with some modifications varying even within one species [30].

**Fig. 1.**
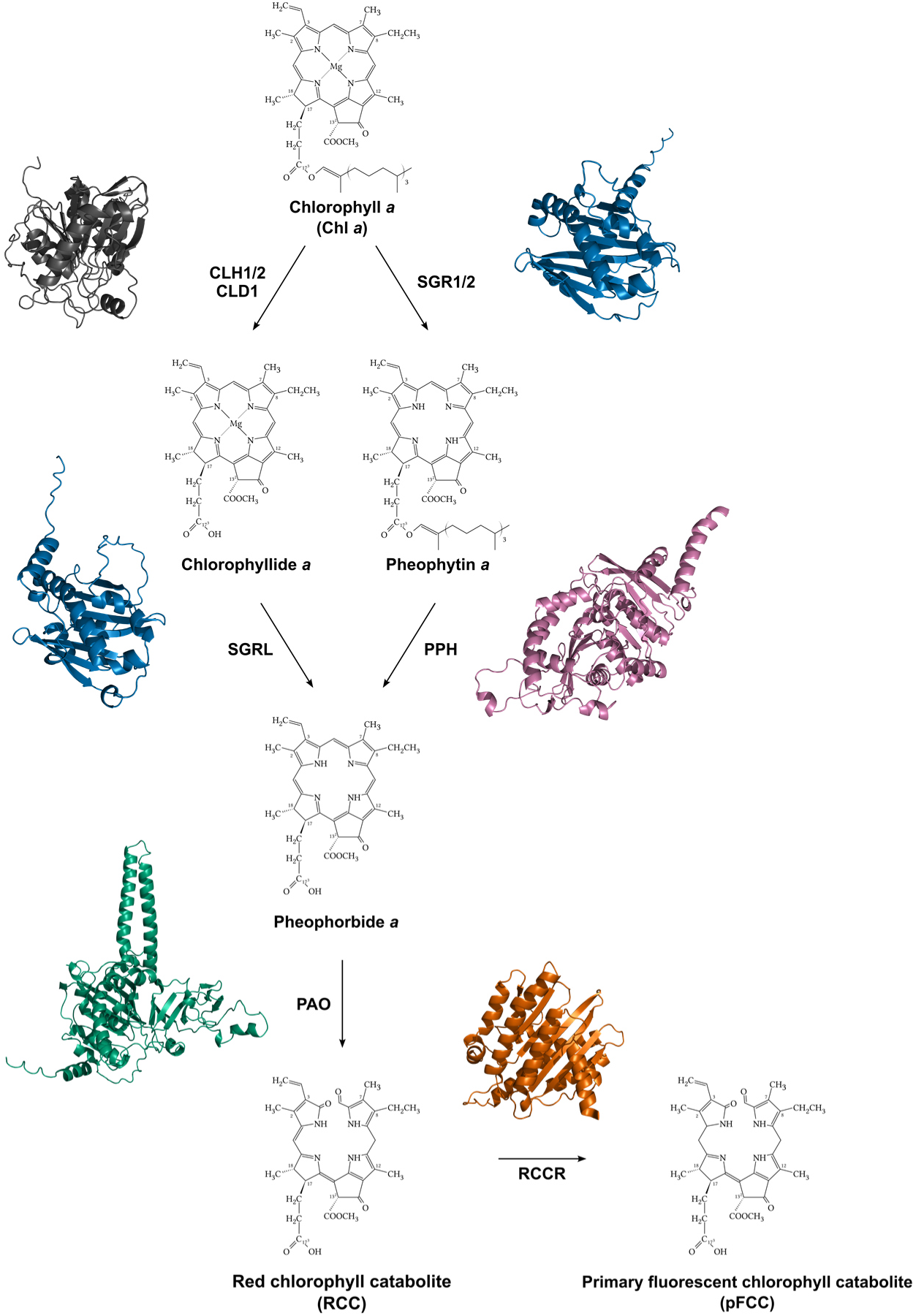
Chlorophyll *a* degradation pathway during leaf senescence in *Arabidopsis thaliana via* the “PAO/phyllobilin” pathway. Starting with Chlorophyll *a (*Chl *a)*, either the magnesium is dechelated first by SGR1 (*STAY-GREEN1)* or SGR2 (*STAY-GREEN2*, shown in vibrant blue) yielding in pheophytin *a*, or the phytol tail is hydrolysed first by CLH1 (*CHLOROPHYLLASE1*, shown in dark grey), CLH2 (*CHLOROPHYLLASE2*) or CLD1 (*CHLOROPHYLL DEPHYTYLASE1*) yielding in chlorophyllide *a*. In the second step, either the magnesium ion is dechelated from chlorophyllide *a* by SGRL (*STAY-GREEN-LIKE*, vibrant blue), or the phytol tail from pheophytin *a* is hydrolysed by PPH (*Pheophytin pheophorbide hydrolase / Pheophytinase*, purple), both yielding in pheophorbide *a*. Next, the macrocycle is opened through a dioxygenation reaction performed by PAO (*pheophorbide a oxygenase*, teal green) yielding in red chlorophyll catabolite, from which The C20-C1 double bond is reduced by RCCR (*red chlorophyll catabolite reductase*, burnt orange) resulting in the formation of primary fluorescent chlorophyll catabolite (pFCC) as terminal product, which is exported from the chloroplast after further modification. The structures of CLH1 (dark grey), SGR2 (blue), SGRL (blue), PPH (purple), PAO (green) and RCCR (orange) were predicted using Alphafold 3 [102] and visualized using PyMOL v3.1 [103].

### Chlorophyll Degradation in other Organisms

Although the Chl *a* degradation pathway is well understood in plants, it remains largely unexplored in other species [31]. Fungi, although not directly studied for Chl *a* degradation, are known for their ability to break down a wide range of complex molecules and polymers [32]. They possess heme oxygenases, which are crucial for obtaining iron from their environment [33] and are capable of opening the tetrapyrrole macrocycle of heme which releases the iron ion and produces biliverdin in one reaction. While both heme and biliverdin are closely related to Chl *a* and its degradation products, whether these or related enzymes can also degrade Chl *a* remains an open question. In multicellular algae SGR homologues were identified across various phyla, which showed that some members of Glaucophyta but none of Rhodophyta possess sequence homologues to the known plant enzymes [34]. In the unicellular algae *Chlamydomonas reinhardtii*, an SGR homologue has been identified, which shows activity towards Chl *a in vitro* but is not involved in Chl *a* degradation during nitrogen starvation [35]. Compared to the observed SGR1 and SGR2 upregulation in *A. thaliana* during senescence [36], this gene is depleted under similar conditions in *C. reinhardtii* [8], again drawing an inconclusive picture. Furthermore, *C. reinhardtii*, but also *Auxenochlorella protothecoide*s, have been reported to excrete RCC-like pigments under stress conditions [37]. Additionally, multiple studies came to the conclusion that RCC formation and its subsequent modifications predate the evolution of multicellularity and land invasion, implying but not proving a fully functional degradation pathway [31, 38, 39]. The knowledge of chlorophyll catabolic enzymes (CCEs) in prokaryotes is similarly thin. Currently only sequence homologues of SGR are reported in a non-photosynthetic member of Chloroflexota [40], as well as in Archaea, but none are identified in Cyanobacteriota [34]. Some of the identified SGR homologues were tested *in vitro*, where their magnesium dechelation activity was found to be highly variable [34]. This led the authors to the conclusion that a protein with promiscuous activity towards Chl *a* has been horizontally transferred from a non-photosynthetic organism to the common ancestor of plants where it gained substrate specificity over time. All other reactions of the Chl *a* degradation pathway are currently unexplored, presenting a huge knowledge gap in one of earths most prevalent processes.

### Beyond sequence – the path towards structure-based homology inference

One reason of Chl *a* degradation being understudied in other organisms is due to the challenge in asserting functional enzyme homologies based on sequences. Over the past decades, protein function inference relied heavily on sequence comparisons, using BLASTP-based sequence similarity search against sequence databases or screening with hidden Markov models (HMMs) profiles derived from multiple sequence alignments (MSAs) of both *in silico* predicted and experimentally characterised sequences [41]. These sequence-based strategies remain relevant, however, are prone to diminished performance in the so-called twilight zone of low sequence conservation [42, 43]. In addition to sequence, the three-dimensional (3D) protein structure has long been recognised to define biological function, and protein structures are often more conserved across evolutionary timescales [44]. However, until recently, structural inference of protein function was challenging due to the lack of available structures as well as algorithms to efficiently compare them, all while the amount of protein sequence data is heavily growing due to advancement in sequencing technologies during the last two decades. Artificial intelligence (AI) based protein structure prediction systems like RoseTTAFold [45], OpenFold [46], and particularly AlphaFold2 (AF2) [47] address these scaling limitations of experimentally determining protein structures, and have paved the way towards mining a previously inaccessible treasure trove of structural data. For instance, Google DeepMind’s and EMBL-EBI’s AlphaFold Protein Structure Database (AFDB) already harboured over 214 million structural predictions of protein sequences one year after the release of AF2. This dwarfs the roughly 180.000 experimentally solved structures over the past 50 years, from which only about 12.000 proteins are unique based on their sequence identity [48, 49]. By using deep learning to predict protein structures from their sequences, these tools also enabled innovation in the field of structural alignment tools like Foldseek [50–52] and Reseek [53], thereby opening a new frontier in homology inference.

In this study, we investigated Chl *a* degradation in prokaryotes, bridging the homology gap across the domains of life by identifying structural homologues of *A. thaliana* chlorophyll catabolic enzymes (CCEs) using Foldseek [50–52] to sieve through the ESM Metagenomic Atlas [54] (https://esmatlas.com/). The resulting structure homologues were then clustered using a structure similarity graph-based technique implemented with NetworkX v3.4 [55] and the Markov Cluster Algorithm (MCL) [56, 57]. Manually curated plant sequence HMMs were then used to identify the functional homologue cluster for each CCE. HMMs from the hereby determined clusters were searched against GenBank to identify prokaryotic isolates containing homologues to all CCEs, resulting in the selection of *Shewanella acanthi* for *in vivo* Chl *a* degradation analysis. In addition, we also examined the global expanse of Chl *a* degradation capability among prokaryotes by analysing publicly available sequencing data, showing its ubiquitous spread among diverse ecosystems and taxa.

## Materials and Methods

### Dataset curation for initial CCE structure search

Chlorophyll catabolic enzymes (CCEs) of *Arabidopsis thaliana* and well-annotated homologues from other plants were retrieved from the UniProt Knowledge Base [58]. To ensure that only true CCEs are included, we selected proteins in the curated Swiss-Prot dataset with an evidence level of either “transcript” or “protein” and a total annotation quality score of at least three. The complete list of the selected proteins and their accession numbers can be found in Supplementary Table S1. If multiple sequences fit our criteria, they were aligned using MAFFT’s L-INS-I algorithm [59] and processed into Hidden Markov Models (HMMs) using the HH-suite3 [60]. If only one protein passed, the cluster verification step (described below) was carried out by aligning the sequence against the cluster HMMs using HMMER3 [61, 62].

### Identification of prokarotic homologues

Protein structures were retrieved from the PDB [63], if available, or predicted using AlphaFold2 [64] or ESM-Fold [65] to balance accuracy and throughput. *A. thaliana* (AT) CCEs were structurally aligned against ∼30 million high-confidence protein structures from the ESM Atlas [65] using Foldseek [66]. As all AT-SGRs returned no hits, we performed a BLASTp search against NCBI’s nr database using their sequences, and predicted the structures of the returned bacterial sequences using ESM-Fold (Supplementary Fig. S2B). After a Foldseek alignment against the AT-SGRs the remaining structures (Supplementary Fig. S2C) were processed into an HMM. For all other CCEs, the structural homologs returned from the ESM Atlas were clustered along a structural similarity graph, where the clustering parameters for the Markov clustering algorithm (MCL) [67] were optimised on clustering modularity. For each CCE the clusters were validated by building hidden Markov models (HMMs) from multiple sequence alignments, which were compared to curated plant HMMs using HH-suite3 [68]. The highest scoring cluster was selected as verified functional CCE cluster for downstream analyses. The workflow is depicted in Figure 2, detailed descriptions can be found in the Supplementary Methods S1, S2 & S3.

**Fig. 2.**
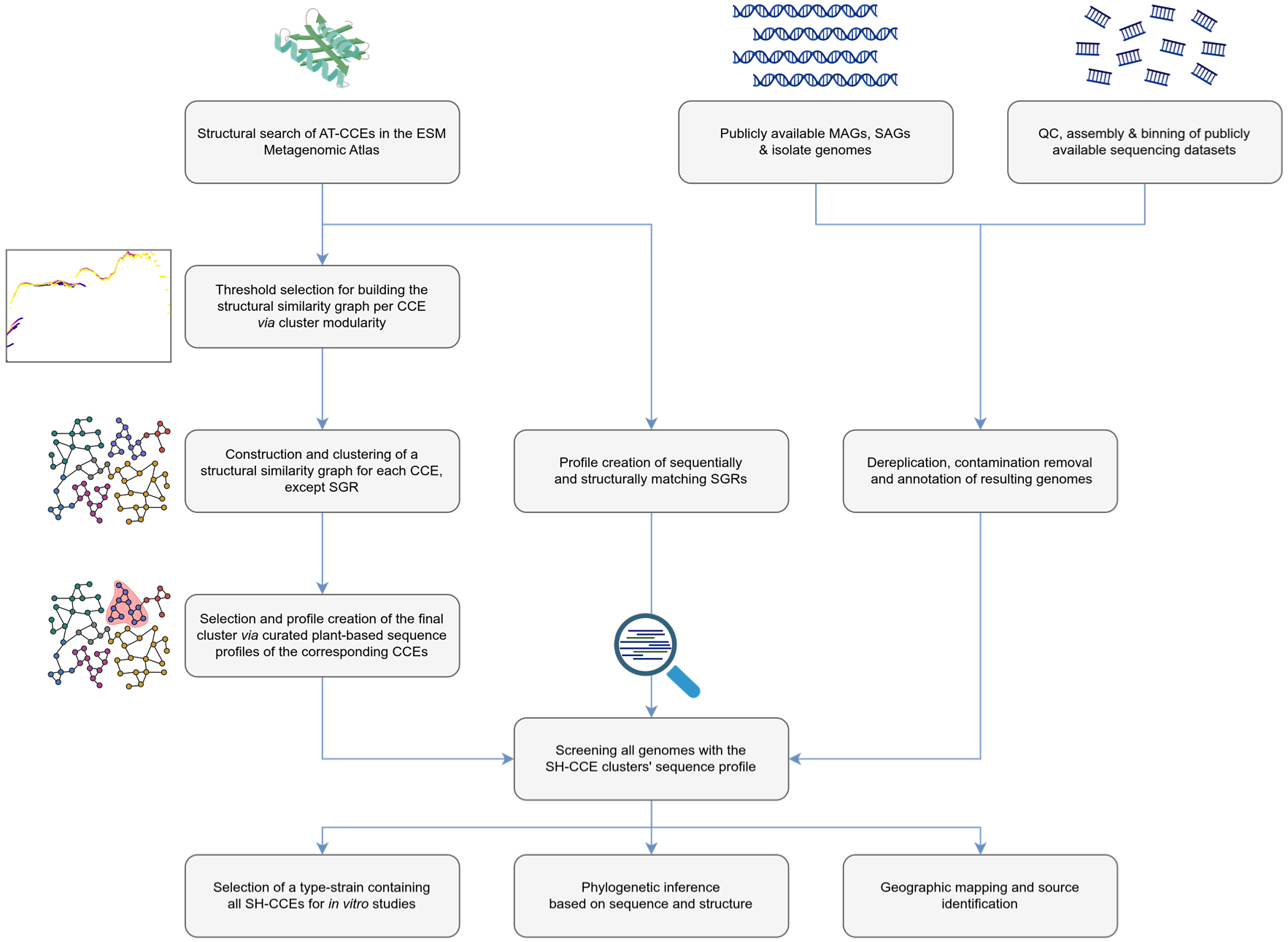
Schematic overview of the bioinformatic framework applied in this study. Structural homologs from *A. thaliana-*CCEs were identified in the ESM Metagenomic Atlas using Foldseek [50, 51]. The resulting structures were then all-vs-all aligned using Foldseek. A structural similarity graph was constructed using NetworkX [55], and clustered with the Markov cluster algorithm (MCL) [104]. After selecting the cluster with the highest profile similarity by searching them against curated plant HMMs using HH-suite3 [60], we obtained one final HMM per *A. thaliana* CCE. In parallel, we prepared publicly available MAGs, SAGs and isolate genomes, as well as own assemblies of MAGs from public metagenomics datasets, and screened them using our SH-CCE HMMs. The resulting alignments were used for phylogenetic and geographic analyses as well as for the selection of a type-strain available from a culture collection to be tested *in vitro*. (Figure was created using icons from BioRender).

### Genomes and strain metadata processing and visualisation

The geographical and environmental distributions of identified prokaryotic genomes containing at least one predicted structural homolog of chlorophyll catabolic enzymes (SH-CCEs) were visualised using metadata extracted from various databases to obtain isolation source and location information. A custom Python script to filter and plot the collected sample metadata on global maps with Cartopy and Matplotlib [69] was used to show the global spread of chlorophyll catabolic capabilities. Another custom Python script was used for microbial source classification, which parses metadata tables and classifies sample origins based on a custom classification scheme. The code applies configurable dictionaries and optional ENVO ontology [70, 71] lookups to generate standardised source categories and detailed classification reports. To identify the strains harbouring the predicted prokaryotic chlorophyll catabolic enzymes (SH-CCEs), we used a custom Python script that integrates taxonomic and strain metadata by matching strain identifiers across multiple reference sources. Specifically, the script reconciles strain information obtained from BacDive (advsearch_bacdive_2025-10-20), the GTDB metadata table (r220), and the NCBI taxdump (downloaded on 2025-10-21) to extract, standardise, and unify all relevant strain-level metadata associated with the input genomes. We performed all metadata visualisation using either custom Python or R scripts, as well as Origin, as indicated in the relevant figure legends. Detailed descriptions of the employed methods for data collection, metagenomic assembly and taxonomic assignment can be found in Supplementary Methods (S4, S5, S6 & S7). Detailed protocols and setups of in vivo experiments, including growth experiments and HPLC analyses can be found ibidem (S8-S12, Supplementary Table S5).

## Results

### Structural alignments using Foldseek

By leveraging well-characterised chlorophyll catabolic enzymes (CCEs) and the degradation pathway in *Arabidopsis thaliana* (AT), we searched for and *in silico* characterised putative prokaryotic CCEs (SH-CCEs). The initial Foldseek search was performed using the MGnify HighConfidence (pLDDT and pTM score ≥0.7) threshold, which resulted in significant hits with all AT homologues except AT-SGR1, AT-SGR2 or AT-SGRL, where only one hit occurred with LowConfidence structures (pLDDT and pTM <0.7). The most predominant matches were observed for SH-PPH, with 69,964 hits, while SH-RCCR returned the fewest, with only 313 sequences. Homologs of SH-PAO and SH-CLH yielded similar numbers of matches, with 7,748 and 9,574 sequences, respectively. In contrast, SH-CLH1 produced nearly double the number of hits compared to SH-CLH2, with 17,962 sequences, and a significant overlap was observed among the hits of the SH-CLH homologs (Supplementary Fig. S2a), leading to the merger of the SH-CLH1 and SH-CLH2 datasets for all following analyses. To enrich our dataset for coverage for SH-SGRs, we performed a BLASTP search against NCBI’s nr database using AT-SGR1, AT-SGR2 and AT-SGRL sequences as queries. The resulting 1522 unique bacterial sequences (*E*-Value ≤1 × 10^−5^) (Supplementary Fig. S2b) were folded using ESM-Fold and the predicted structures were aligned against the AT-SGRs using Foldseek. We obtained 58 unique structures (*E*-value ≥1 × 10^−5^ and a target coverage >80 %) (Supplementary Fig. S2c).

### Structural similarity-based graph clustering and sequence-based hidden Markov model verification

To define the bitscore limit and inflation for the graph-based clustering, we first obtained the all-vs-all structural alignment bitscores per CCE using Foldseek, and explored the modularity distribution across them by exhaustively iterating through them as described in Supplementary Method S1-3 and is depicted in Fig. S3 A-D). The selected parameters for each CCE, listed in Supplementary Table S1, were applied to generate the final clusters along with the MSAs and HMMs of each cluster, resulting in one final graph representation per AT-CCE (Supplementary Fig. S3 E-H). The “verified cluster” was then determined by aligning the cluster HMMs against the corresponding curated plant HMM using the HH-suite3, and selecting the highest scoring cluster’s HMM for all downstream analyses.

### Prediction of prokaryotic CCEs in global sequencing datasets

To retrieve SH-CCEs, we analysed over a billion proteins across at least 352,847 reported medium- to high-quality metagenomes, including MAGs and SAGs, and isolated genomes from various projects. Based on our high-confidence SH-CCEs HMMs, initial screening of the predicted proteomes identified 161,619 genomes harbouring at least one SH-CCE homolog. To normalize genome quality, we excluded 1,509 previously reported medium- to high-quality genomes from various studies that did not meet our criteria of at least 50% completeness and less than 10% contamination, estimated by CheckM2 [72]. The final dataset comprises 160,110 genomes containing over 480 million predicted proteins (Supplementary Table S3, Supplementary method S4-S6). Our analysis of global (meta)-genomic data revealed a highly diverse landscape of putative SH-CCEs, highlighting the potential role of prokaryotes in the degradation of Chl *a* (Fig. 3). We examined a total of 160,110 genomes, which exhibited a nearly ubiquitous distribution across the Earth’s biosphere (Fig. 3A). The distribution of these genomes varied significantly by environmental, with about 31% of the genomes encountered in marine samples. Additionally, a substantial number of genomes were derived from human sources (Host-Associated-human: ∼20%) and, to a lesser extent, from fresh water sources (∼15%), reflecting the diverse ecological roles of prokaryotes in sequestering Chl *a* (Fig. 3A). Interestingly, only about 1.8% of the identified SH-CCEs have been linked to plant-associated sources (Fig. 3B), which include plant parts such as root, nodules, and leaves or plant-linked biomes such as the phyllosphere and rhizosphere.

**Fig. 3.**
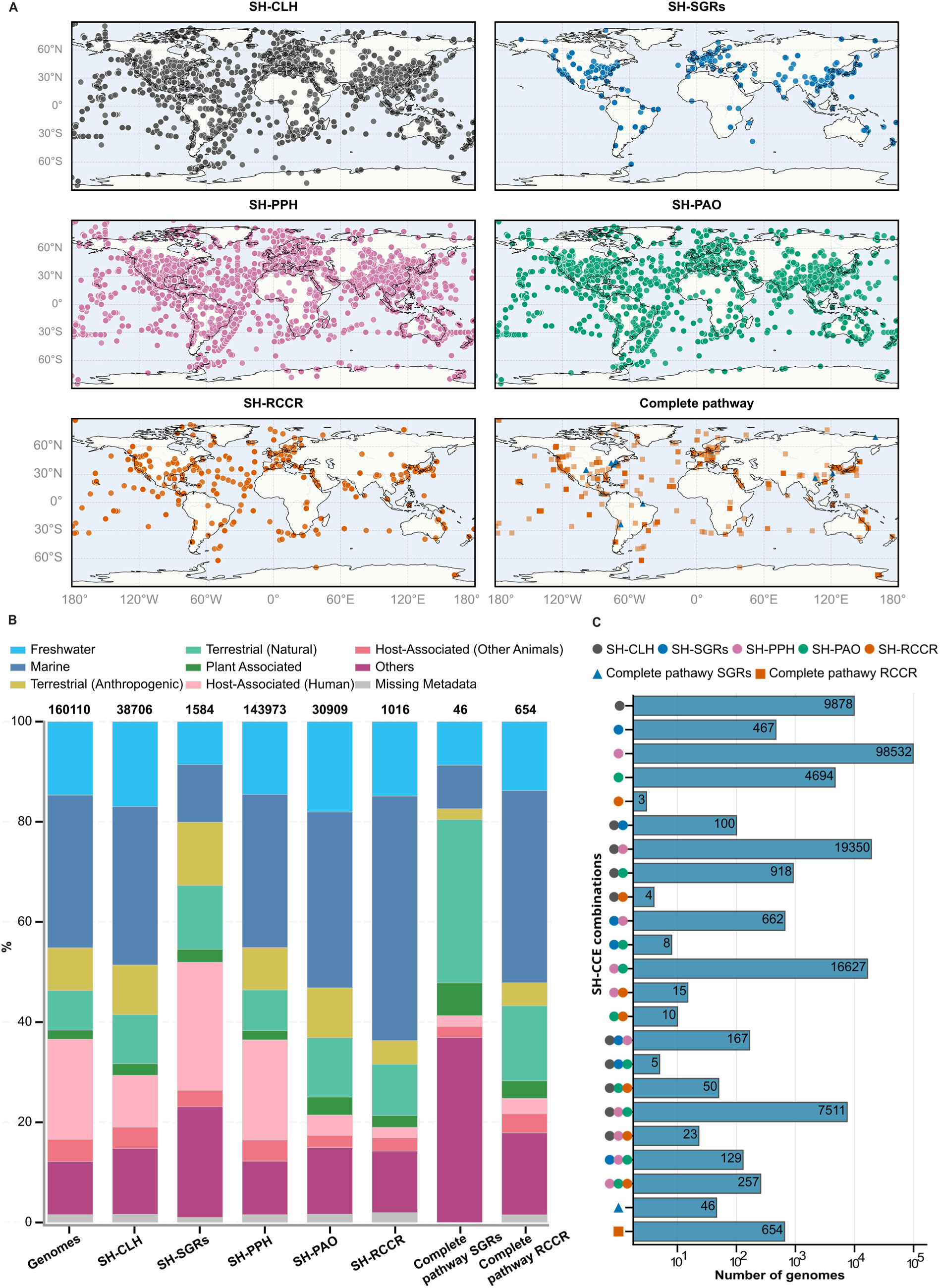
Global distribution of prokaryotic genomes harbouring structural homologs of CCEs. **A**: global map showing the spatial distribution of genomes harbouring SH-CLH (dark grey), SH-SGRs (blue), SH-PPH (purple), SH-PAO (green), and SH-RCCR (orange). **B**: bar chart illustrating the habitat-level representation of prokaryotic genomes that encode structural homologues of CCEs. Each column shows the proportion of genomes (first column) or proportion of genomes encoding the putative enzymes per habitat type with colours corresponding to individual habitat types, with the total genome number for each column represented in bold font (top). **C**: bar plot showing the number of prokaryotic genomes harbouring one or more CCE structural homologue generated *via* computing the binary presence (1) or absence (0) of proteins across the dataset and subsequent translation into protein combination. The bars display the frequency of genomes containing specific combinations of SH-CCEs. We generated the global map using Cartopy for geographic projections, with Matplotlib [69] handling data visualisation.

Of the predicted 480 million proteins, we identified 431,332 putative SH-CCEs among the various medium- to high-quality genomes (Supplementary Table S3), of which approximately 73% are SH-PPHs. Notably, 164,768 of these proteins (∼52%) were the sole putative SH-CCEs identified in 98,532 genomes (∼62%) (Fig. 3C). Conversely, we identified 1,193 (∼0.3%) and 1,674 (∼0.4%) putative SH-RCCR and SH-SGRs homologs, respectively, among the analysed genomes. Unlike the widespread SH-PPH homologues, only three genomes harbour SH-RCCR as singletons and occur sparingly in combination with any one other SH-CCEs (Fig. 3C). Interestingly, the presence of SH-RCCR and SH-SGRs homologs in any genome appears to be mutually exclusive. Therefore, genomes carrying the three other SH-CCEs (SH-CHL, SH-PAO and SH-PPH) alongside SH-RCCR or SH-SGRs are considered here to constitute two separate Chl *a* degrading pathways. However, absence of a key SH-CCE in an otherwise complete pathway may suggest a Chl *a* degradation bottleneck. Previous studies have reported the secretion of RCC in heterotrophic green alga *Auxenochlorella* that lacks RCCR homologs [73, 74]. By contrast, *Chlamydomonas* SGR mutants show no Chl *a* degradation disruption, suggesting the presence of alternate Mg-dechelatases [8]. In the present context, the occurrence of SH-SGRs or SH-RCCR alongside SH-CHL, SH-PPH and SH-PAO indicates a complete Chl *a* degradation pathway, here termed the complete pathway SGRs or RCCR, respectively (Fig. 3B-C). Despite the predominance of the SH-CCEs in the marine samples, only the complete pathway RCCR (∼38%) follows this trend, with the natural terrestrial environment contributing more genomes (∼33%) with the complete pathway SGRs (Fig. 3B). In the lower end of the range, host-associated (human), plant-associated, and anthropogenic terrestrial sources each accounted for only about 3-4% of complete pathway RCCR SH-CCEs. However, when considering the relative abundance of genomes within the isolation sources, plant-associated and natural terrestrial sources contained a higher percentage of complete pathway RCCR SH-CCEs, comprising ∼0.8% of the genomes, compared to only ∼0.5% in the marine environment. Additionally, plant-associated sources represented a more significant proportion of “complete pathway SGRs” SH-CCEs, with 0.1% of their genomes, whereas marine-associated genomes contained just 0.008% of these elements.

To estimate the diversity of taxonomically distinct genomes with putative SH-CCEs, we deduplicated and decontaminated the genomes, resulting in 74,734 genomes representing distinct species across 147 prokaryotic phyla (137 bacterial and 11 archaeal). These include seventy-nine named phyla and sixty-eight with taxonomic placeholders, according to the GTDB scheme. Fig. 4 depicts the named phylawhile the extended version comprising all identified phyla (including taxonomic placeholders) is listed in Supplemental Material (Supplementary Fig. S4 & Supplementary Table S4). Among the bacterial genomes, members of five phyla *Pseudomonadota* (∼29%), *Bacteroidota* (∼16%), *Bacillota* (∼13%), *Actinomycetota* (∼12%), and *Chloroflexota* (∼5%) account for over 70% of the catalog of SH-CCE-harboring genomes (Fig. 4). Collectively, these five phyla, along with Cyanobacteriota, comprise ∼ 99% of the genomes containing a complete set of the four SH-CCEs (including either SH-RCCR or SH-SGR). *Actinomycetota* predominates, contributing 156 (∼33%) SH-RCCR-type genomes, followed closely by *Pseudomonadota* and Cyanobacteriota, each representing ∼24% of the SH-RCCR-type genomes. In contrast, members of Bacillota account for 31 of the 33 SH-SGR-type genomes, with no SH-RCCR-type genomes observed within this phylum. Although no genome, and by extension, no phylum, contains a complete combining SH-RCCR and SH-SGR homologs type, we identified one SH-SGRs-type genome (GTDB placeholder: sp016929795) within *Chloroflexota*. Although several archaeal genomes harbour large numbers SH-CCEs, only one member placed at order level (*Sigynarchaeales*, family: SOKP01) showed the full complement of the putative enzymes albeit with an SH-SGR (Fig. 4). Further analysis of the bacterial genomes with potential for complete Chl *a* degradation revealed that 262 (56% of the total) have cultured representation, distributed among thirty bacterial families (Supplementary Fig. S5). Among these, members of the families *Nostocaceae*, *Micromonosporaceae, Mycobacteriaceae, Streptosporangiaceae, and Nocardioidaceae* are particularly dominant, collectively accounting for 68% of the isolates (Supplementary Fig. S5), highlighting the ecological role and functional specialization of these families. Interestingly, a substantial proportion of the genomes (158) have isolates that are traceable to various culture collections (Supplementary Fig. 5), providing a valuable resource for experimental validation and further biochemical characterization. While SH-CCE identification and their taxonomic and spatial distributions are described above, detailed phylogenetic analyses and interpretation based on 3Di and AA characters are provided in Supplementary Method S7, Supplementary Note S1 and S2.

**Fig. 4.**
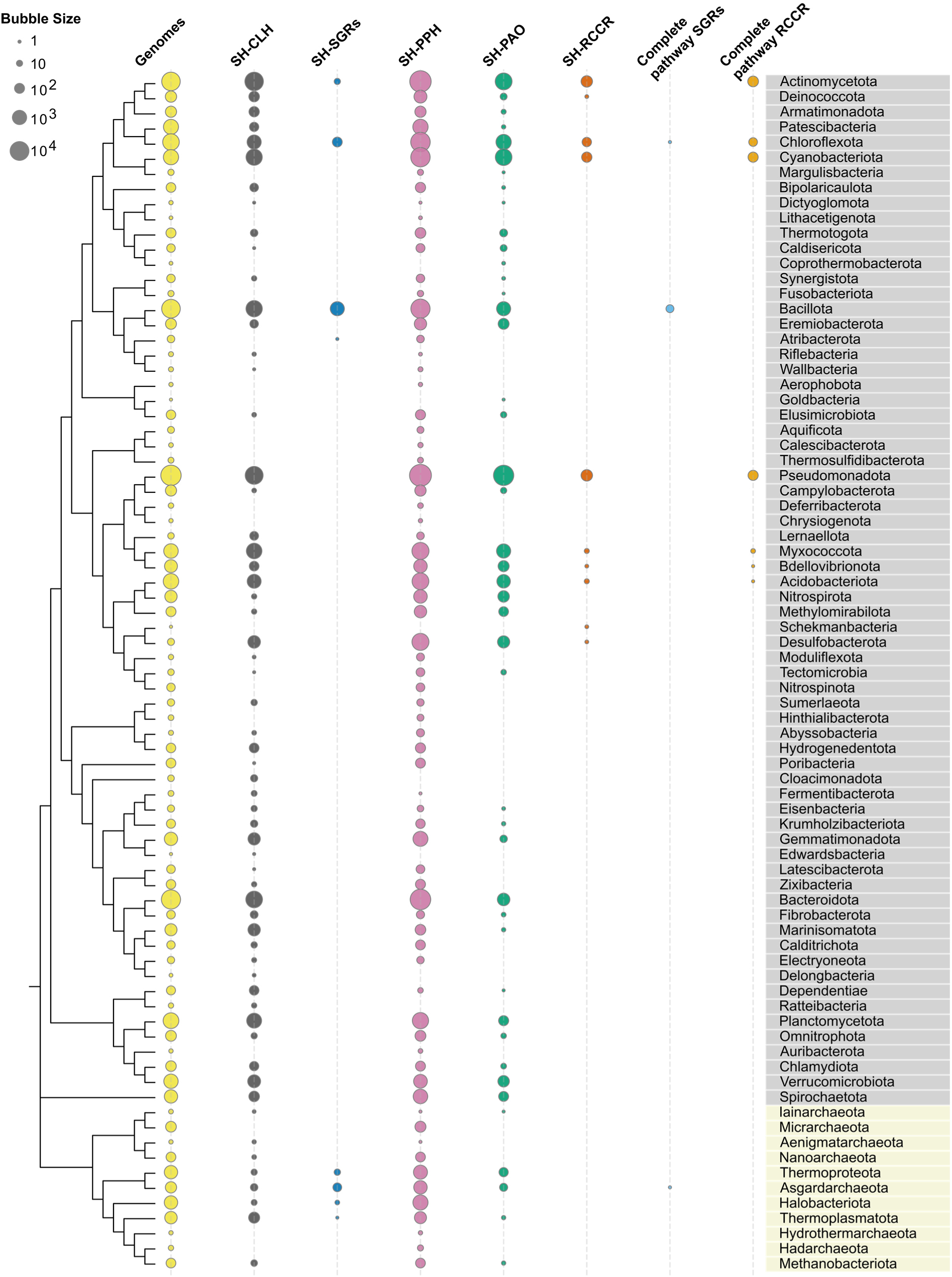
Distribution of predicted structural homologs of CCEs among seventy-nine named prokaryotic phyla, comprising 74,230 distinct species. An additional 504 species affiliated to distinct but unnamed phyla are included in Supplementary Fig. 4. The midpoint-rooted phylogenetic tree (only topology displayed: Left), based on the concatenated and trimmed core proteins of 79 representative genomes (5,776 sites), was built using IQ-TREE with the model LG+I+G4 and ultrafast bootstrap approximation. Bacterial phyla are highlighted in light gray, while archaeal phyla are in light beige. The bubble plots are generated with Matplotlib [69] and correspond to the number of distinct SH-CCEs predicted in each phylum. The bubble sizes are scaled logarithmically.

### Chlorophyll *a* degradation by *Shewanella acanthi*

To determine whether the *in silico* predictions translate into an *in vivo* function we conducted degradation experiments using the *S. acanthi* wild-type (WT) and the ΔPPH-ΔCLH mutant in artificial seawater (ASW) supplemented with 10% chlorophyll extract from spirulina powder. Figure 5 depicts a representative experiment run, showing the bottles with medium just before inoculation (Fig. 5A) and after two days of incubation in the dark at 22°C and 200 rpm of shaking (Fig. 5B). The WT samples show a clear discolouration compared to the ΔPPH-ΔCLH mutant and the sterile control.

**Fig. 5.**
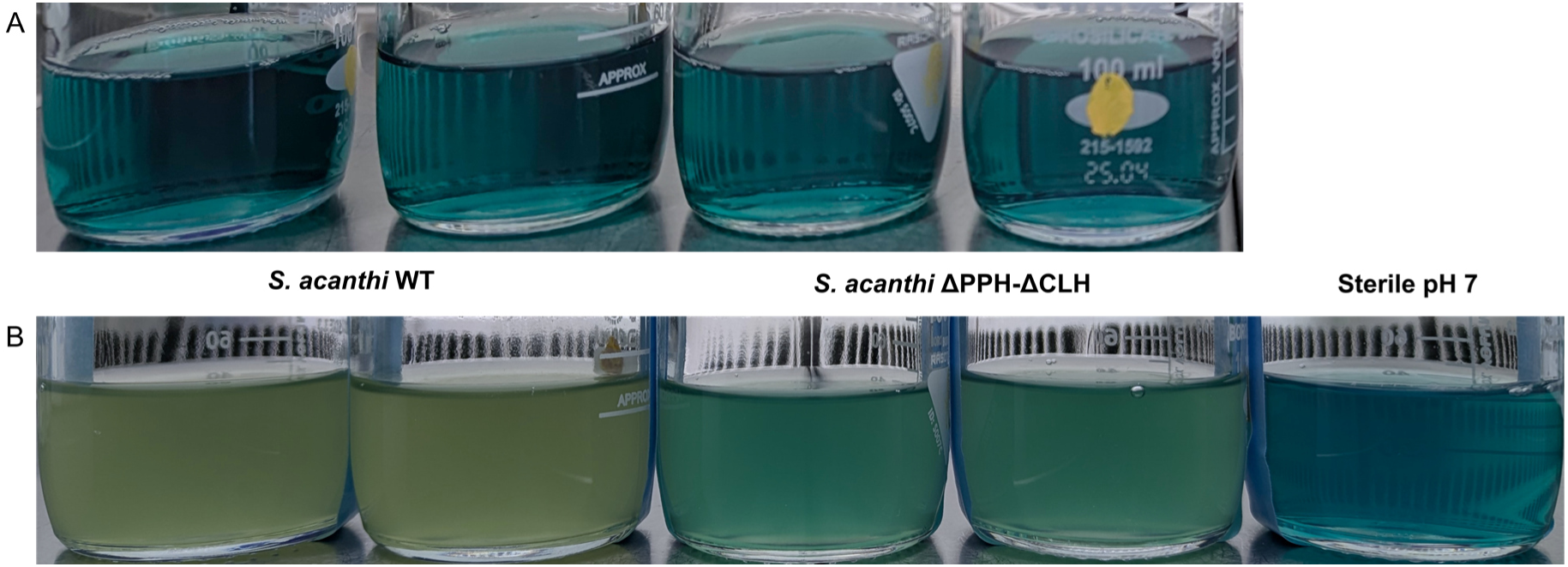
Exemplary images from the *in vivo* chlorophyll degradation experiments conducted with the *Shewanella acanthi* wild-type (WT) and the *Shewanella acanthi* ΔPPH-ΔCLH mutant, showing a clear discolouration of the medium by the WT compared to the mutant. **A:** Just before inoculation to an OD600 of 0.04. **B:** Two days after incubation at 22°C and 200 rpm shaking, average OD600 *S. acanthi* WT: 0.485, average OD600 *S. acanthi* ΔPPH-ΔCLH: 0.445

Fluorescence spectra were obtained directly after inoculation, from where on all inoculated samples, as well as the sterile control at approximately pH 4.5, showed a shift of the initial main peak at ∼660 nm towards ∼685 nm (Fig. 6A, B and C). In the inoculated samples the intensity at ∼685 nm was the lowest at T6 from where on it increased again for the remainder of the experiment (Fig. 6A and B). In the sterile control at approximately pH 7, the fluorescence intensity showed a steady decrease between 600 nm and 700 nm with only a slight shift of the peak from 660 nm to ∼665 nm and no additional peaks forming (Fig. 6C). The sterile control at approximately pH 4.5 also showed a gradual shift towards 685 nm, while retaining a secondary peak at ∼650 nm, with the overall intensity decreasing over the whole course of the experiment (Fig. 6C). Since spirulina extract was used in the experiments and not pure Chl *a,*due to its poor solubility in water, HPLC analyses were performed to reveal the actual degradation profiles (Fig. 6D and Supplementary Method S12). We analysed samples taken at T0, T6, T8, T18 and T25 from inoculated samples, while only T0, T8 and T25 were analysed for the sterile controls. The data shows the conversion of Chl *a* to Pheo *a* in all samples, except for the sterile control at approximately pH 7, where the Pheo *a* concentration varied only slightly across all sampling points (Fig. 6D). The sterile control at approximately pH 4.5 showed the fastest conversion to Pheo *a*, with a large part of the Chl *a* already converted before the first sample was taken (Fig. 6D). The WT followed a similar trajectory, albeit the initial conversion proceeding slower, with both WT and the sterile pH 4.5 sample showing a slight accumulation of Phebide *a* over time, and the levels decreasing again between the last two WT samples (Fig. 6D and Supplementary Fig. S6). Meanwhile the ΔPPH-ΔCLH mutant showed an almost linear decease in Chl *a* concentration, coupled with a gradually increasing Pheo *a* and constant low Phebide *a* concentration (Fig. 6D and Supplementary Fig. S6). Chlorophyllide *a* was non-detectable in all samples (Supplementary Method S12).

**Fig. 6.**
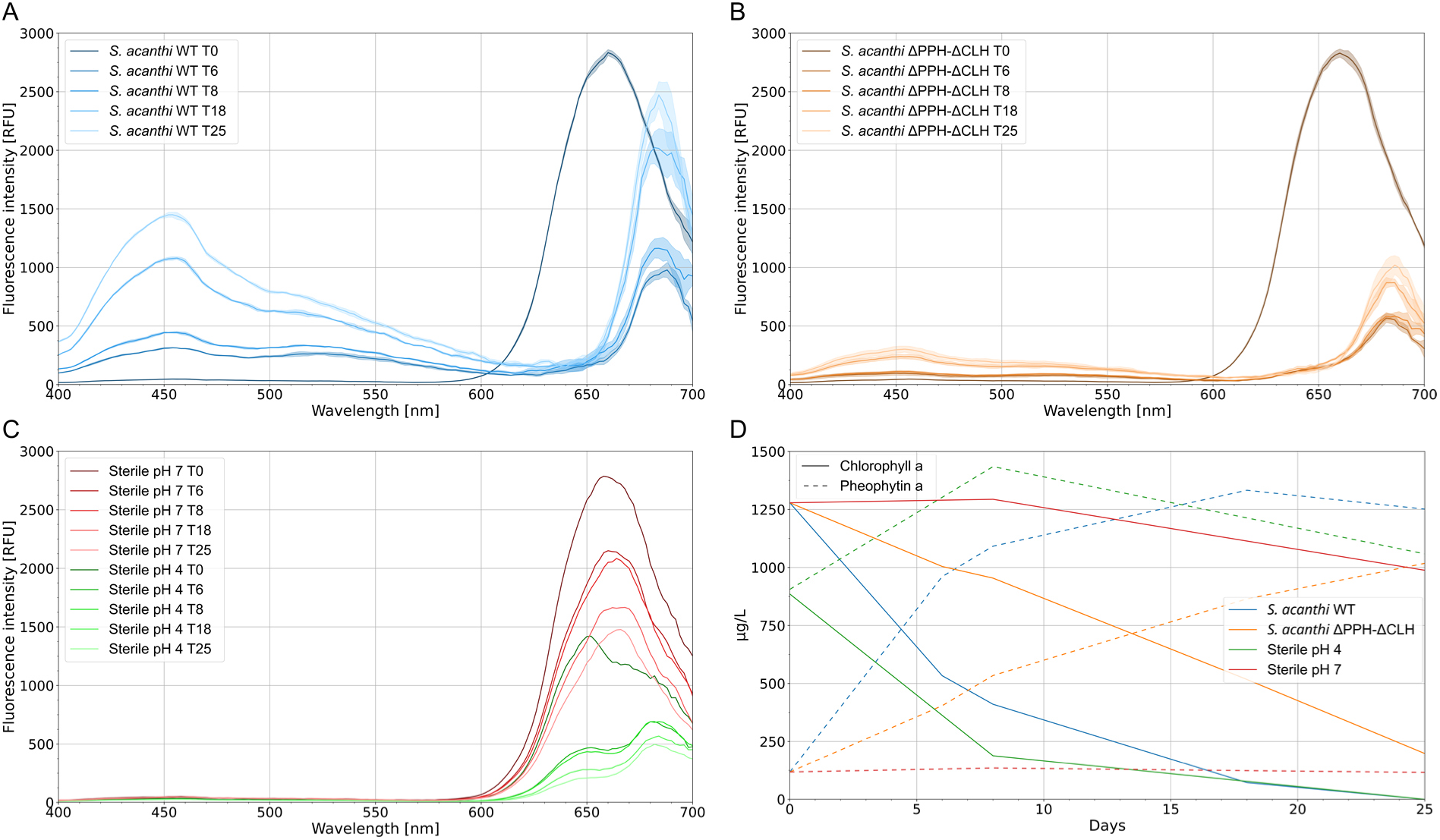
Fluorescence spectra (λex 365 nm) recorded during the degradation experiment at the time of inoculation (T0), as well as 6, 8, 18 and 25 days after inoculation (T6, T8, T18, T25). The spectra were collected as 100 µl samples of biological duplicates from **A:** *Shewanella acanthi* WT, **B:** *Shewanella acanthi* ΔPPH-ΔCLH, **C:** sterile controls at pH 7 and pH 4. For biological samples, each line represents the average of duplicates, with the variance between replicates shown as opaque area. As a post-processing step, they were smoothed using a sliding window with a size of three, where the average was applied to the first value in the window. The HPLC-based monitoring of Chl *a* degradation is shown in **D:** Chl a (solid), Pheo a (dashed).

## Discussion

Recognising novel or rare biological functions across different evolutionary lineages remains a significant challenge in modern biology, often due to the challenge in asserting functional enzyme homologies based on sequences. Information about protein structure can provide an alternative and robust approach, enabling the elucidation of shared functional insights that may be obscured at the sequence level. Here, we leverage recent advances in structural analysis to identify prokaryotes capable of breaking down chlorophyll (Chl) *a,* which is arguably one of the most abundant pigments on earth. Since the degradation pathway is best described in *Arabidopsis thaliana*, we used its proteins as structural references to identify and compare them with their prokaryotic functional relatives using a custom workflow.

Clustering distant sequences poses a significant computational challenge, as traditional alignment techniques often struggle to identify weak sequence-level relationships. Among various methods, the fast and scalable unsupervised clustering algorithm MCL effectively approaches this challenge. It leverages graph-based clustering to group sequences based on connectivity in a similarity network rather than direct similarity [75, 76]. Because it relies on similarity scores, clustering similar structures using MCL may lead to saturation and poor graph resolution, because every structure has some alignment score with all other proteins in the pool. To improve cluster detection and focus on stronger relationships within the clusters, we introduced a bitscore limit favouring smaller clusters of high structural similarity instead of clustering to the general core fold. Notably, various factors, including the length of the proteins, influence the bitscore; therefore, limits must be determined individually for each enzyme. The inflation parameter of the MCL algorithm affects the granularity of clustering; a higher inflation value favours smaller clusters, while a lower value results in fewer, larger clusters [76]. In determining the inflation value for the MCL algorithm, we found that the lowest tested value, 1.2, was consistently preferred. The preference for lower values likely stems from the fact that the bitscore limit had already separated some clusters, meaning that a higher inflation value would have resulted in over-clustering of the graph. While clustering algorithms like MCL organise data into meaningful groups, the reliability and biological relevance of these clusters depend heavily on effective cluster verification. Extrinsic measures that rely on some external ground truth can provide an unbiased means of assessing cluster validity; however, often no prior information exists, imposing a reliance solely on the network’s internal structure [77]. Therefore, we optimised the clustering modularity, since it is a performant, intrinsic clustering evaluation metric of the NetworkX module. Based on the resulting modularity, bitscore limits, and inflation plots, we aimed to prioritise the earliest possible local maximum, assuming that excluding weaker edges would disrupt inter-cluster connections as little as possible. For the SH-PAO dataset, the parameters were adjusted to around the inflection point of the modularity increase at a bitscore limit of 840, since this configuration yielded significantly better results during the HMM verification compared to the local maximum at a bitscore limit of 450. To verify our predicted homologues, we employed strict selection criteria. In addition to the protein membership in Swiss-Prot, the curated plant enzymes must have had a UniProtKB annotation score of at least three and a minimum of mRNA experimental evidence. These stringent requirements limited sequence options to those conferring the desired function, reducing diversity in the multiple sequence alignment (MSA) and, ultimately, in the generated hidden Markov model (HMM). To enhance the detection of conserved domains within these diverse sequences, we utilised MAFFT’s high-accuracy local alignment algorithm, L-INS-i [78]. This method is known for its high accuracy, although it is less efficient when handling large datasets. Because of this limitation, we aligned only the curated set of plant enzymes and the clusters of SH-RCCR using MAFFT. For all other SH-CCEs, we opted for MAGUS [79], which employs a “divide and conquer” approach, allowing for the alignment of large sequence clusters while maintaining the high accuracy characteristic of MAFFT’s local alignment. After generating the MSAs, we processed them into HMMs using HH-suite3 [60] by combining information about the position-specific conservation of amino acids with substitution matrices to estimate the evolutionary cost of residue changes and insertions and deletions at specific positions within the MSA [80, 81]. The HMM-HMM alignments between the HMMs created for each cluster of SH-CCE and the corresponding curated plant CCE HMMs provide greater sensitivity than through HMM sequence alignments alone [60].

The enzymes that catalyse phytol hydrolysis, specifically AT-CLH1, AT-CLH2, and AT-PPH, yielded the most hits with structural homologues in prokaryotes, likely due to their more universal enzyme function. All three belong to the large and diverse family of α/β hydrolases, however, despite their diverse enzymatic activities, they share relatively low sequence identities, but a highly conserved [82]. This led to hits within the MGnify HighConfidence database and a considerable overlap between the returned proteins. In contrast, no suitable targets were found for the AT-SGRs, even though prokaryotic functional homologues of these Mg-dechelatases have been reported [34, 40, 83]. This suggests their under-representation in MGnify or a generally lower distribution among bacteria. Similarly, the low number of hits returned for AT-PAO and AT-RCCR might be due to the limited size of the MGnify structures database, which consists of only 30,157,582 structures [84].

Integrating the highlighted structure-based screening and the high purity of our final prokaryotic proteome database provided a comprehensive and novel approach for high confidence and accurate identification of prokaryotes with the potential capability to degrade Chl *a*. In contrast to previous studies based on similarity searches, our findings suggest a wide distribution of the homologues of CCEs among prokaryotes. For instance, previous reports suggest the absence of its PPH orthologs among the *Cyanobacteriota* using blastP-based screening [13, 74]. However, our structure-based screening of the well-curated dataset revealed 1,678 distinct species affiliated to *Cyanobacteriota* coding at least one putative SH-PPH that co-occurs with a minimum of one other SH-CCE. Instructively, we have predicted various combinations of the SH-CCEs in 74,735 distinct species across 148 prokaryotic phyla. Since the oxidative cleavage of pheophorbide *a* by PAO represents a specific reaction in Chl *a* degradation [9], yielding the highly phototoxic RCC [24], we considered the presence of SH-PAO in combination with the majority of the other enzymes (SH-CHL, SH-PPH, SH-RCCR and SH-SGRs), as a primary marker for the presence the pathway or part thereof in combination with the majority of the other enzymes. Two classes of prokaryotes harbouring complete or near full complement of SH-CCEs, named the SH-RCCR and SH-SGR types were found. The latter SH-CCE type mainly occurs among the predominantly plant-associated *Bacillota* and cannot remove RCC. The absence of the putative SH-RCCR will ordinarily preclude the likelihood of the function in the SH-SGR type organisms. However, previous reports of RCC in heterotrophic green alga lacking RCCR homologs [73, 74] suggest an ecological role for this partial chlorophyll degradation. Sequestration of the toxic RCC presumably occurs in a community that includes other chlorophyll degraders, such as the SH-RCCR-type class, and possibly others with a functional SH-RCCR. Unlike the SH-SGRs-type, SH-RCCR-types potentially could complete the breakdown of Chl *a*, despite the missing SH-SGRs due to the promiscuous activity of the Mg-dechelatases as demonstrated with *Chlamydomonas* SGR mutants [8].

We further found the complete Chl *a* degradation pathway among photoautotrophic *Cyanobacteriota* and *Pseudomonadota*, with the former utilising the light-harvesting complexes along with the pigments recycling mechanisms like plants [85, 86]. However, the predominant taxa predicted to harbour the complete Chl *a* degradation pathway occur among the heterotrophic *Actinomycetota* and *Pseudomonadota*, constituting ∼55% of the 469 identified species. The heterotrophic members of these phyla play a central role in deconstructing complex molecules in various biomes [87–90]. The environmental and taxonomic distribution of the predicted Chl *a*-degrading prokaryotes indicates the widespread nature of this function and the ecological role of the organisms, despite possible biases towards certain taxa in public databases.

### *In vivo* studies

To test the ability of prokaryotes to deconstruct chlorophyll *a* (Chl *a*), we selected the chemoheterotrophic bacterium *Shewanella acanthi* (KCTC 82650) as a model organism based on two major factors. Firstly, *Shewanella acanthi* encodes the full SH-RCCR-type genotype based on the *in silico* analysis, and all top-scoring SH-CCE hits reside within an approximately 11 kbp region. Secondly, the genus *Shewanella* typically possesses favourable culturing conditions, as well as established genetic systems for some members [91, 92]. As a long standing model organism for studying extracellular electron transport [93–95], it therefore provided a good candidate model to investigate the Chl *a* degradation pathway *in vivo*.

The first step in the degradation of Chl *a* involves magnesium de-chelation catalysed by SGR. This process can also occur non-enzymatically under acidic conditions [96, 97], while the following phytol hydrolysis catalysed by CLH or PPH could also be performed by non-specific hydrolases, although with presumably way lower reaction rates, with both leading to an initial discolouration of the pigment. This aligns with our observations over the course of the degradation experiments, where no discolouration was observed for the sterile controls at approximately pH 7, with an immediate discolouration of the sterile controls at approximately pH 4.5 (Fig 6). The discolorations didn’t progress further, compared to the biological samples containing either the *S. acanthi* WT or the *S. acanthi* ΔPPH-ΔCLH mutant, where strong and fast colour changes were observed for the WT, while the ΔPPH-ΔCLH mutant showed a drastically slower change. The intermediate colours align with previous descriptions of pheophytin *a* spectra [98, 99] Notably, a high ratio of pheophytin *a* to Chl *a* has been reported in coastal regions, reaching up to 0.48 in Massachusetts Bay [100] and 0.7 in the Mediterranean Sea [101], supporting the prevalence of prokaryote-mediated Chl *a* degradation in the natural environment.

Since not pure Chl *a* was used for the experiments due to poor solubility but spirulina extract, HPLC analyses was performed in addition to reveal the Chl *a* degradation profiles. The data was consistent with the fluorescence measurements, corroborating a clear Chl *a* degradation as well as Pheo *a* formation.

## Conclusion

Our findings reveal a previously overlooked aspect of chlorophyll-related chemistry in prokaryotes. By using plant chlorophyll catabolism enzyme (CCE) as reference points, we identified previously unknown distant prokaryotic homologs. Recent advances in computational methods, particularly high-confidence AI structural predictions and comparative genomics, confirm the presence and activity of these pathways in prokaryotes. Combining these computational approaches with experimental feedback loops will enhance the discovery and provide a clearer understanding of the cryptic ecological functions in prokaryotes

## Availability of data and materials

All data generated or analysed during this study is included in the main article and its supplementary materials. Metagenome-assembled genomes (MAGs) have been deposited in Zenodo (https://doi.org/10.5281/zenodo.17827782). Custom scripts used for data analysis are also available at Zenodo (https://doi.org/10.5281/zenodo.18027694). Supplementary tables and associated metadata have also been archived at Zenodo (https://doi.org/10.5281/zenodo.18028059). All resources will be publicly accessible upon publication.

## Acknowledgements

We thank David Thiele for support with the laboratory work, and John Vollmers for bioinformatics support.

## Authors contributions

H.A., H.F., G.S. and A.K.K. wrote the manuscript, H.A. and H.F did the data curation and designed and performed the computational analyses, H.F. and G.S. performed the *in vivo* studies and A.K.K conceived the study.

