## Supporting Information for "Chlorophyll a degradation in Prokaryotes"

Supplementary Method S1. Protein structure prediction and alignment

To predict the structures of proteins AlphaFold2 [1] and ESM-Fold [2] were used, depending on the size of the dataset in question, to balance the trade-off between AlphaFold2's high resource and time usage and the accompanying gain in accuracy, with ESM-Fold's high throughput and its slightly lower accuracy. An overview of the bioinformatic workflow is depicted in Fig. 2. All *A. thaliana* (AT) CCEs were folded using AlphaFold2 [1], except for AT-RCCR, where the published crystal structure (PDB ID:2ZXL) [3] was selected. We used the high-quality (pLDDT and pTM score ≥ 0.7) structure predictions from the ESM Metagenomic Atlas (ESM HighConfidence) [2] as search database, amounting to 30.157.582 high quality predicted protein structures. Foldseek [4] was used to structurally align the AT-CCEs against this database, using the 3Di+AA alignment mode, an *E*-value threshold of $1\times10^{-5}$ and a target coverage threshold of 80% to include target structures matching only one domain of the query and to account for transit peptides in the AT-CCEs. As AT-SGR1, AT-SGR2 and AT-SGRL returned no hits within ESM HighConfidence and only one hit in the full dataset, we enriched our dataset by performing a BLASTP search against NCBI's nr database using their sequences as queries. The resulting bacterial sequences (*E*-Value $\leq 1\times10^{-5}$) (Supplementary Fig. S2b) were dereplicated and folded using ESM-Fold. The predicted structures were then aligned against the AT-SGRs using Foldseek with the same thresholds as for the other CCEs (*E*-value $\geq 1 \times 10^{-5}$ and a target coverage >80 %) resulting in 58 unique structures (Supplementary Fig. S2c), which were treated as a "verified cluster" and processed into a HMM, as described for the other CCEs after structural similarity-based graph clustering.

Supplementary Method S2. Structural similarity-based graph clustering to identify functional structure clusters

To identify the subset of proteins which are the presumable functional CCE homologues among the structural homologues identified in the ESM HighConfidence database we devised a structural similarity-based graph clustering workflow. To this end NetworkX [5] was used to create an undirected graph for each CCE from the structural alignment scores obtained by self-aligning the initially identified structural CCE homologues without any thresholds and Foldseek's pre-filter disabled. On the resulting graph, the individual proteins are represented by nodes, connected *via* edges, with the alignment bitscore assigned as weight. Two nodes are only connected with an edge if the alignment bitscore exceeds an exhaustively determined bitscore limit between the two proteins. This threshold, as well as the inflation parameter required by the Markov Clustering Algorithm (MCL) [6], was individually determined for every protein by optimising the parameters to maximise the modularity Q, defined and simplified in equations 1 & 2 [7,8], of the clustering. Modularity measures how well a graph is divided into clusters by comparing the number of connections within clusters and across clusters to a theoretical null model, i.e. what a random clustering would look like. A higher modularity indicates stronger, more clearly defined clusters, but simply optimizing for the maximum modularity, does not guarantee to result in the clustering which represents the underlying reality [9].

$$Q=\frac{1}{2m}\sum_{ij}\left(A_{ij}-\gamma\frac{k_i k_j}{2m}\right)\delta\left(c_i,c_j\right) \qquad (1)$$

$$Q=\sum_{c=1}^{n}\left[\frac{L_c}{m}-\gamma\left(\frac{k_c}{2m}\right)^2\right] \qquad (2)$$

To determine the lowest bitscore limit producing a local modularity maximum, it was calculated for every bitscore limit and inflation value in a given range. To determine the bitscore limit and inflation parameters to use, we exhaustively iterated through increasing bitscore limits in coarse step sizes of 10 and with a starting inflation parameter of 1.4 for MCL. After a local modularity maximum was identified by plotting the bitscore limits, the inflation and the resulting modularity using Plotly [10], a refinement iteration was started with bitscore limit step sizes of 5 or 1, depending on the size of the dataset, in the range around the prior identified maximum. The inflation was simultaneously varied from 1.2 to 2.1, except for the SH-CLH and SH-PPH datasets, where the inflation only varied up to 1.6 due to the large dataset size, and since it became increasingly clear that lower inflation values were favourable (Supplementary Table S2 & Supplementary Fig. S3). The hereby determined parameters were applied to the final clustering. For the graph visualisation, the nodes are distributed based on the Fruchterman-Reingold force-directed algorithm [11], which solves the graph structure by treating edges between nodes as springs, with the weight of the edge as the pulling force, while nodes repel each other. By iteratively simulating the forces between the nodes, they are moved on the graph until their positions are close to an equilibrium.

<u>Supplementary Method S3. Sequence-based hidden Markov model verification of structure clusters</u>

To obtain an HMM for each cluster on the graph, we aligned clusters with fewer than 500 members using MAFFT's high accuracy L-INS-I algorithm [12]. For larger clusters we used MAGUS [13], which enables efficient computation of MSAs from large protein sets by splitting the dataset according to a guide-tree, whose leaves are individually aligned using MAFFTs L-INS-I algorithm, and then joined into one large MSA [12]. The HMMs generated from the cluster's MSA were combined into one HMM database per CCE using the HH-suite3 [14] and subsequently aligned against the curated plant-based HMMs, where the cluster with the highest HMM alignment score is selected as a "verified cluster" for all subsequent analyses. The MSAs of all verified clusters were finally processed into HMMs using HMMER3 [15,16] for HMM-sequence alignment against the various (meta-)genomic datasets.

<u>Supplementary Method S4. Genome collection for identifying chlorophyll catabolic enzymes (CCEs) in prokaryotes</u>

To determine the distribution of the predicted prokaryotic structural homologues (SH) of Chl *a* degradation proteins, we retrieved 352.847 medium- to high-quality genomes with over 50% completeness and less than 10% contamination estimates, which spanned different habitats across the Earth's biosphere. The genomes comprised 52.515 MAGs from the Earth's

Microbiomes (GEM) catalogue [17], 23.755 MAGs, 5.160 SAGs, and 1.697 reference genomes from the Ocean Microbiomics Database (OMD) [18–21], 122.865 genomes from the Global Microbial Gene Catalogue (GMGCv1) [22], 112.022 genomes from the GTDB-r220 database [23] and 12.670 MAGs reported in one of our previous studies [24]. Furthermore, 904 metagenome samples were selected based on pre-screening of the 308.6 million proteins in the Global Ocean Gene catalogue 1.0 [25]. 289 additional marine and soil metagenomic samples were retrieved from the European Nucleotide Archive (ENA; www.ebi.ac.uk/ena), assembled and binned resulting in an additional 19.405 and 2.758 medium- to high-quality MAGs (for procedure see Supplementary Method S5).

Supplementary Method S5. Metagenome assembly, binning and taxonomic assignment

In addition to the genomes, MAGs and SAGs obtained from the databases and projects outlined above, we downloaded 1,193 raw read samples from ENA, as well as the metadata of all datasets. Raw reads were trimmed using fastp v0.23.4 [26] and assembled using MEGAHIT v1.2.9 [27] with the preset “meta-large” and a minimum contig size of 1000. Assembled metagenomes were binned using Aviary v0.9.0 [28], which uses multiple binning algorithms to generate high-quality bins. For genome quality evaluation, the generated bins and retrieved genomes were subsequently evaluated with CheckM2 [29]. All genomes were finally taxonomically annotated with reference to GTDB release 220 using GTDB-Tk v2.4.0 [23,30].

Supplementary Method S6: Identification of prokaryotic chlorophyll catabolic enzymes (CCEs)

To identify prokaryotic CCEs, a total of 352.847 medium to high-quality genomes were annotated using PRODIGAL v2.6.3 [31]. The HMMs generated from each verified structural homology CCE (SH-CCE) cluster were searched against the predicted proteins using HMMER3 [15,16]. Hits with a full sequence *E*-value below 1e-14 and a score ≥ 70 were considered positive for SH-CCEs detection. Genomes harbouring CCEs were dereplicated with galah 0.4.0 [32] cluster mode, and CheckM2’s [29] quality report was used for genome filtering and ranking. To improve the prediction accuracy of SH-CCEs for the final set of 74,734 non-redundant genomes, we filtered the genomes using MDMcleaner v0.8.7 [33]. We identified 39, 245 genomes (~53%) that contained between 0.002% and 20% of sequences flagged as “fraction_delete”, indicating definite contaminating fractions (Supplementary Table S4). These contaminating sequences (~89,691 contigs) were removed prior to downstream analysis.

Supplementary Method S7: Phylogenetic reconstructions using amino acids and 3Di sequences

To determine the evolutionary relationships among the prokaryotic, as well as between them and the plant homologues, we used a custom Python script to filter the top-scoring predicted prokaryotic chlorophyll catabolic enzymes (SH-CCEs) in each genome containing the predicted complete SH-CCE sets. Overall, the selected prokaryotic homologues comprise the highest-confidence predicted SH-CCEs per genome, including 469 sequences each of SH-CLH, SH-PPH and SH-PAO, 436 sequences of SH-RCCR, and 33 sequences of SH-SGRs. For each set, we selected the corresponding plant homologues, including eight homologues (CLH2_ARATH, CLH1_ARATH, PAO_ARATH, PPH_ARATH, RCCR_ARATH, SGRL_ARATH, SGR2_ARATH and SGR1_ARATH) of *Arabidopsis thaliana*, 5 (PAO_ORYSJ, Q69XR3_ORYSJ, RCCR1_ORYSJ, SGR_ORYSJ and SGRL_ORYSJ) of *Oryza sativa* subsp. *japonica*, 2 (RCCR_HORVU and F2E504_HORVV) of *Hordeum vulgare* and one sequence each of *Capsicum annuum* (SGR_CAPAN), *Chenopodium album* (CLH0_CHEAL), *Citrus* *sinensis* (CLH1_CITSI), *Litchi chinensis* (A0A0E3KDE4_LITCN) and *Lolium perenne* (A0A0U3ABU7_LOLPR). The 3D structures of the 1.896 prokaryotic and plant CCEs were predicted using AlphaFold3 [34]. The resulting structures for each homologue set were used to generate alignments with FoldMason [35]. Multiple sequence alignment and refinement was performed with MAFFT v7.525 [12,36]. The amino acid (AA) and tertiary-interaction characters (3Di) alignments were then trimmed using trimAl v1.4.rev15 [37] and BMGE v1.12 with BLOSUM30 as substitution matrix [38]. For AA phylogenies, we reconstructed maximum likelihood (ML) trees using IQ-TREE v3.0.1 [39] with extended model selection (MFP) followed by inference, 1000 ultrafast bootstrap replicates, 1000 replicates of the Shimodaira-Hasegawa approximate likelihood ratio test [40], and optimised UFBoot trees by NNI on the bootstrap alignment. Using IQ-TREE v3.0.1, ML inferences for 3Di alignments were first performed with MFP and subsequently with the "-mset" option using using the three available 3Di substitution models models Q.3Di.AF, Q.3Di.LLM, 3DiPhy [41,42], as well as the PMB and LG models. Resulting phylogenies were rooted using the minimal ancestor deviation (MAD) method [43] and a custom R script was used to collapse all monophyletic branches and to generate annotations for the final tree visualisation in iTOL [44].

Supplementary Method S8: Growth of *Shewanella acanthi*

A lyophilised culture of *S. acanthii* was purchased from the Korean Collection for Type Cultures (KCTC). The culture was resuspended using 200 µL of LB and incubated at room temperature for 20 minutes. The rehydrated cells were transferred into a culture tube containing 5 mL of LB and also streaked onto corresponding agar plates using an inoculation loop. The liquid cultures and agar plates were incubated overnight at 30°C, with the liquid culture being shaken at 180 rpm. Single colonies were obtained, and glycerol cryostocks were prepared from one of the colonies and stored at -80°C, while the colony's identity was verified *via* 16S rRNA colony PCR coupled with Sanger-sequencing. To prepare the cultures used for the degradation experiments, LB medium was inoculated using the verified cryostocks and incubated over night at 30°C and 200 rpm.

The ΔPPH-ΔCLH mutant of *S. acanthi* was obtained *via* a targeted genome editing service at CreativeBiogene (USA). The mutant was generated by homology directed deletion using a chloramphenicol-sacB based suicide vector system, and verified by us *via* shotgun sequencing on a NextSeq1000 (Illumina, USA), with the reads subsequently mapped against the reference genome of *S. acanthi* to check for the correct deletion of the desired genes. Growth conditions were the same as for the WT.

Supplementary Method S9: Spirulina Extract Preparation

Chl *a* is highly hydrophobic and therefore proved difficult to solve in an aqueous medium. This would have made homogenous sampling during the experiments impossible without the addition of polar solvents which pose a high risk of cytotoxicity or growth inhibition. Therefore, we opted to use a spirulina-based extract for the degradation experiments. It was prepared from spirulina powder obtained at Aldi-Süd (Germany) and extracted by adding 35 ml of $ddH_2O$ to 0.6 g of spirulina powder, vortexing until the powder was fully suspended, and centrifugation at 14.000 x g for 45 min. Afterwards the supernatant was discarded and the washing was repeated with 35 ml $ddH_2O$, and centrifugation at 14.000 x g for 30 min, after which the supernatant was discarded again. Finally, 35 ml $ddH_2O$ and 35 µl Tween20 (Roth, Germany) were added to the pellet, and the suspension was thoroughly vortexed and incubated at 4°C overnight. The next day, the suspension was centrifuged again at 14.000 x g for 45 min, the supernatant was kept and sterilised by filtration through 0.22 µm filters.

Supplementary Method S10: Chlorophyll Degradation Experiments with *S. acanthi* WT & *S. acanthi* ΔPPH-ΔCLH

20 mL pre-cultures of *S. acanthi* WT as well as *S. acanthi* ΔPPH-ΔCLH were prepared in LB and incubated overnight. After sub-culturing them to an $OD_{600}$ of 0.2 they were harvested between an $OD_{600}$ of 0.8-1.0. The cultures were washed thrice by centrifugation at 8,000 g and 4°C for 5 minutes. The supernatant was discarded, and the cells were resuspended in 10 ml of fresh pre-cooled artificial sea water (ASW), and kept on ice until inoculation. Chlorophyll degradation experiments were conducted in duplicates of both strains, as well as single sterile controls at a pH ~7 and pH ~4 in borosilicate screw-neck bottles with their lids not fully tightened down, to allow for gas exchange during cultivation. 50 ml ASW (Supplementary Table S5), supplemented with 10% spirulina extract was seeded with the washed pre-cultures to an $OD_{600}$ of 0.05 and incubated in complete darkness at 22°C and 200 rpm. 100 µl samples were taken directly after inoculation (T0), as well as after 6, 8, 18, and 25 days (T6, T8, T18, T25), into 96 well plates with a UV-transparent bottom (Corning, USA) for spectral analysis with a BioTek Synergy H1 microplate reader (Agilent Technologies, USA). HPLC samples were taken for T0, T6 and T8 time points and frozen until further processing.

Supplementary Method S11: Spectral analysis of Chlorophyll Degradation using a Microplate Reader

Absorbance and fluorescence spectra were measured from the 100 µl samples taken during growth experiments by monitoring emission wavelengths specific to Chl *a* and its degradation products. Fluorescence spectra were recorded at an excitation wavelength of 365 nm and a set gain of 100, from 400 nm to 700 nm in 2 nm steps, with four measurements averaged per emission wavelength. After collection, a custom python script was used to calculate the average of the replicates, and the fluorescence intensities were smoothed using a sliding window approach.

Supplementary Method S12: HPLC analyses

HPLC analyses were carried out by CreativeProteomics (USA), using a Supelcosil Column with a flow rate of 1.0 mL/min and 70:30 (v/v) methanol, 28 mM aqueous TBAA, pH 6.5 for the mobile phase A and methanol for mobile phase B. The percentage of mobile phase was linear from 5% to 100% in 20 min. Detailed sample preparation protocols provided by the company, along with the corresponding HPLC chromatograms, are available on Zenodo (https://doi.org/10.5281/zenodo.19064797).

### Supporting Information Data

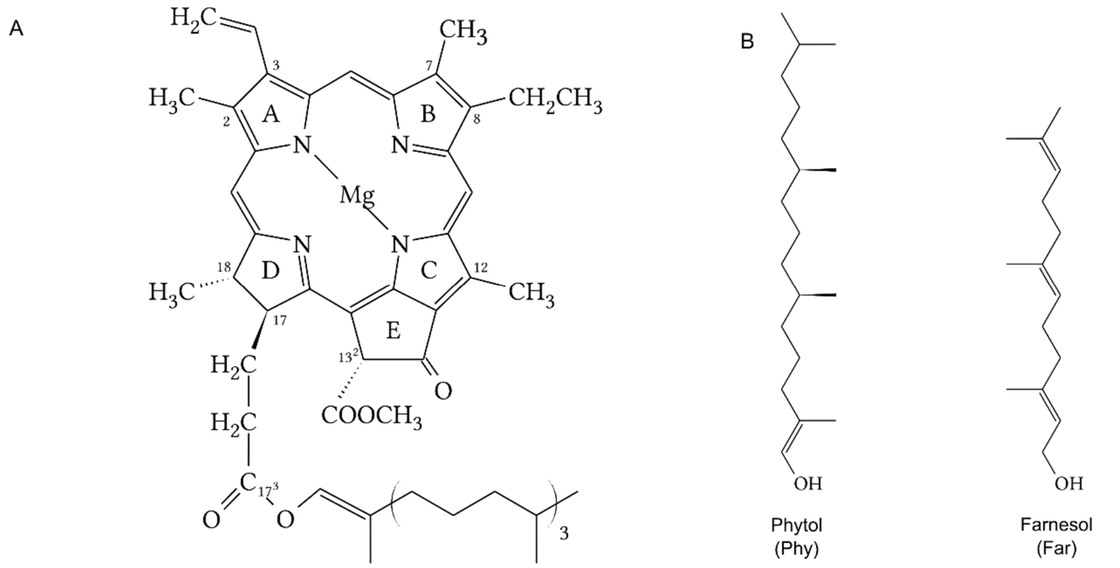


| Name | C3 | C7 | C8 | 7,8-bond | C12 | C13 2 | C20 | C17 3 | 17,18bond |
|---|---|---|---|---|---|---|---|---|---|
| Chl a | -CH=CH2 | Me | Et | double | Me | -COO-CH3 | H | Phy | single |
| Chl b | -CH=CH2 | -CHO | Et | double | Me | -COO-CH3 | H | Phy | single |
| Chl c1 | -CH=CH2 | Me | Et | double | Me | -COO-CH3 | H | H | double |
| BChl a | -CO-CH3 | Me | Et | single | Me | -COO-CH3 | H | Phy | single |
| BChl b | -CO-CH3 | Me | =CH-CH3 | single | Me | -COO-CH3 | H | Phy | single |
| BChl c | -CHOH-CH | Me | Et,Pr,iB | double | Me, Et | H | Me | Far | single |

**Supplementary Fig. S1**. Chlorophyll structures and their side chain modifications. **A**: Chlorophyll *a*, base macrocycle of chlorophyll *a*, important carbon atoms and rings of the macrocycle are labelled after the IUPAC-IUB nomenclature [45], **B**: phytol and farnesol tails; Me = methyl-group, Et = ethyl-group, Pr = n-propyl-group, iB = isobutyl-group.

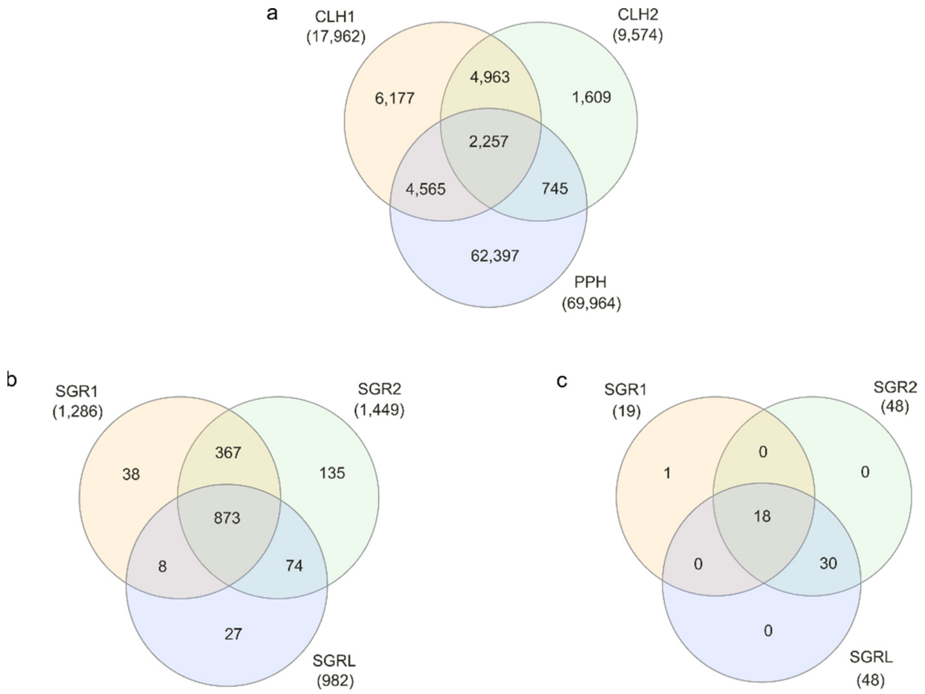


**Supplementary Fig. S2**. Foldseek search against the AT-CCE and SH-SGR enrichment, created using InteractiVenn [46]. **A**: overlap between the Foldseek search results for the phytol-hydrolases AT-CLH1, AT-CLH2 and AT-PPH, **B**: BLASTP search results against NCBI's nr database, constrained to the bacterial domain, with an E-value ≤1 × $10^{-5}$ using the sequences of AT-SGR1, AT-SGR2 and AT-SGRL as queries. **C**: Foldseek search results against the SH-SGR structures obtained from NCBI and predicted by ESM-Fold, with an E-value ≤1 × $10^{-5}$, using the structures of AT-SGR1, AT-SGR2 and AT-SGRL as queries.

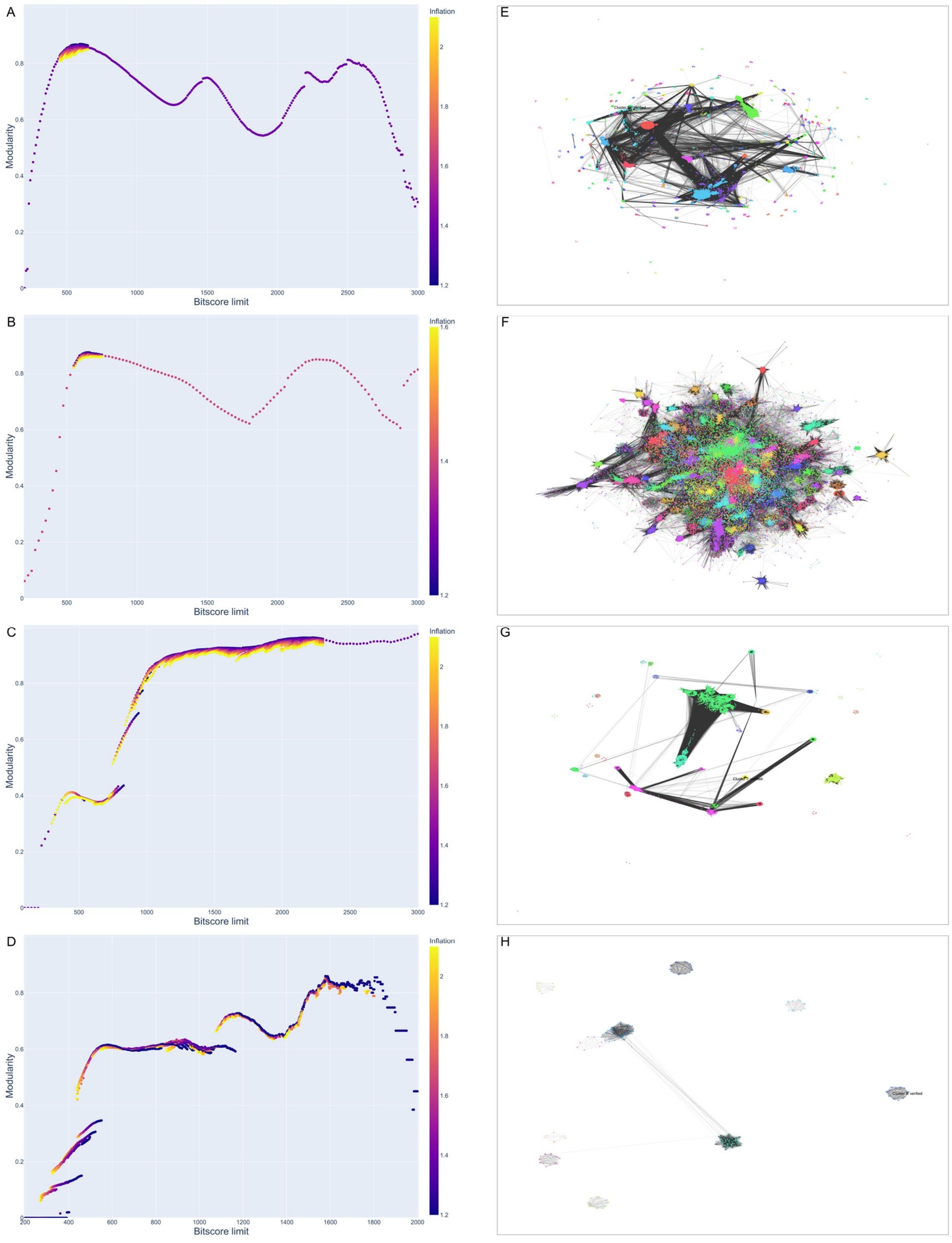


**Supplementary Fig. S3: A-D**: Bitscore limit vs modularity and inflation plots used to determine the final graph clustering parameters. The modularities were calculated by exhaustively iterating through bitscore limits, with a step size of 5 or 10, depending on the dataset size, for graph construction, and varying MCL's inflation parameter from 1.2 to 2.1 or 1.6, again depending on the dataset size, in steps of 0.1, to cluster the structural similarity graph. **E-H**: Structural similarity graphs for each of the CCEs except for the SGRs, since they were handled separately due to low representation in the ESM Metagenomic Atlas. The graphs are constructed from Foldseek structural alignment bitscores using NetworkX and clustered by MCL with the previously determined bitscore limit and inflation parameters. The selected SH-CCE cluster with the highest sequence profile similarity to the corresponding curated plant-CCE HMM is indicated on the graph. **A, E**: SH-CLH, **B, F**: SH-PPH, **C, G**: SH-PAO, **D, H**: SH-RCCR.

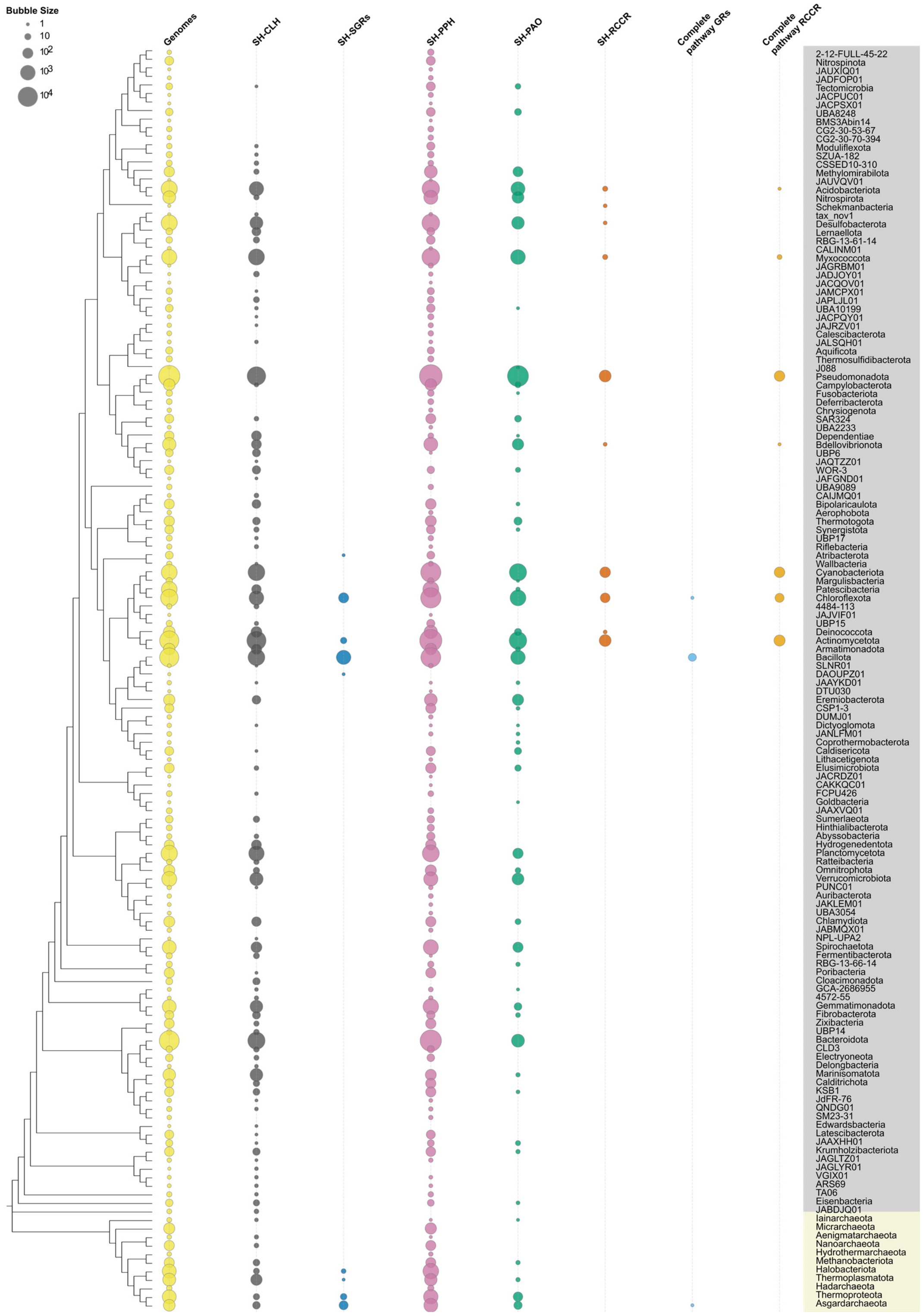


**Supplementary Fig. S4**. Distribution of predicted structural homologs of CCEs among 148 prokaryotic phyla, comprising 74,734 distinct species. The midpoint-rooted phylogenetic tree (only topology displayed: left) based on the concatenated and trimmed core proteins of 148 representative genomes (4,040 sites) built using IQ-TREE with the model LG+I+G4 and ultrafast bootstrap approximation. Bacterial phyla are highlighted in light cyan, while archaeal phyla are in light grey. The bubble plots are generated with Matplotlib and correspond to the number of distinct SH-CCEs predicted in each phylum. The bubble sizes are scaled logarithmically to highlight differences across a wide range of values. Smaller values are enlarged for better visibility, while larger values are proportionally represented. This approach ensures that the scaling emphasizes relative differences while maintaining a visually intuitive representation of the data.

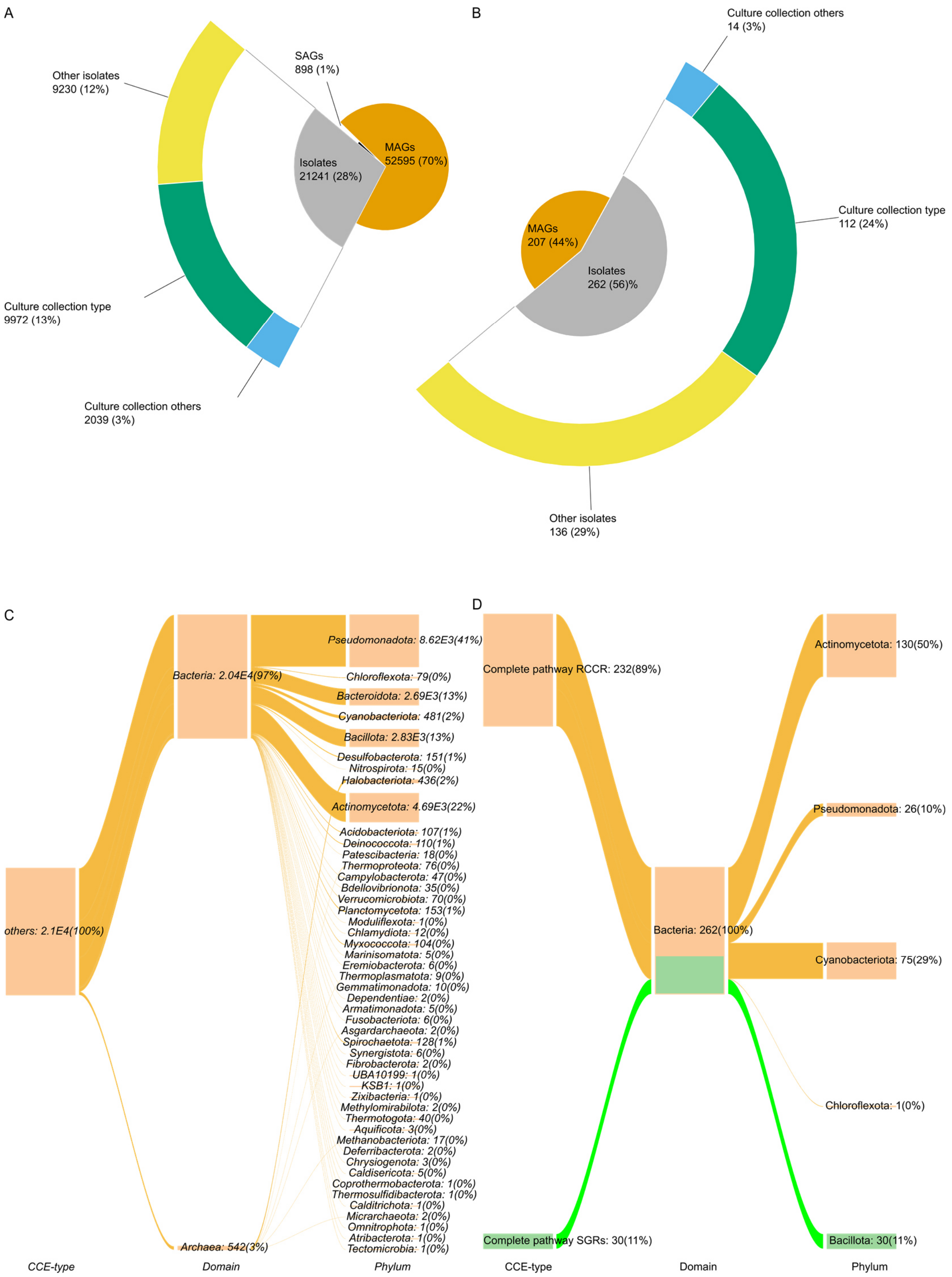


**Supplementary Fig. S5**. Availability of strains that harbour homologs of the predicted prokaryotic chlorophyl catabolic enzymes (SH-CCEs). **A**: Overview of isolate genomes, metagenome-assembled genomes (MAGs) and single amplified genomes (SAGs) among the final 74,734 non-redundant (representative) genomes reported in the current work. We also highlight the proportion of genomes with a record of the corresponding strains in various culture collections. **B**. proportions of the genome types among 469 species-level representative genomes putatively possessing a complete Chl *a* degradation pathway, including the fractions of the corresponding strains available in various culture collections. **C**. Phylum-level affiliation of 20,979 isolates harbouring partial Chl *a* degradation pathway. Phylum-level affiliation of the 262 isolates with documented isolation records and harbouring the complete Chl *a* degradation pathway. Strain and genome metadata were generated based on the harmonised information obtained from BacDive (advsearch_bacdive_2025-10-20), the GTDB metadata table (r220), and the NCBI taxdump (downloaded on 2025-10-21).

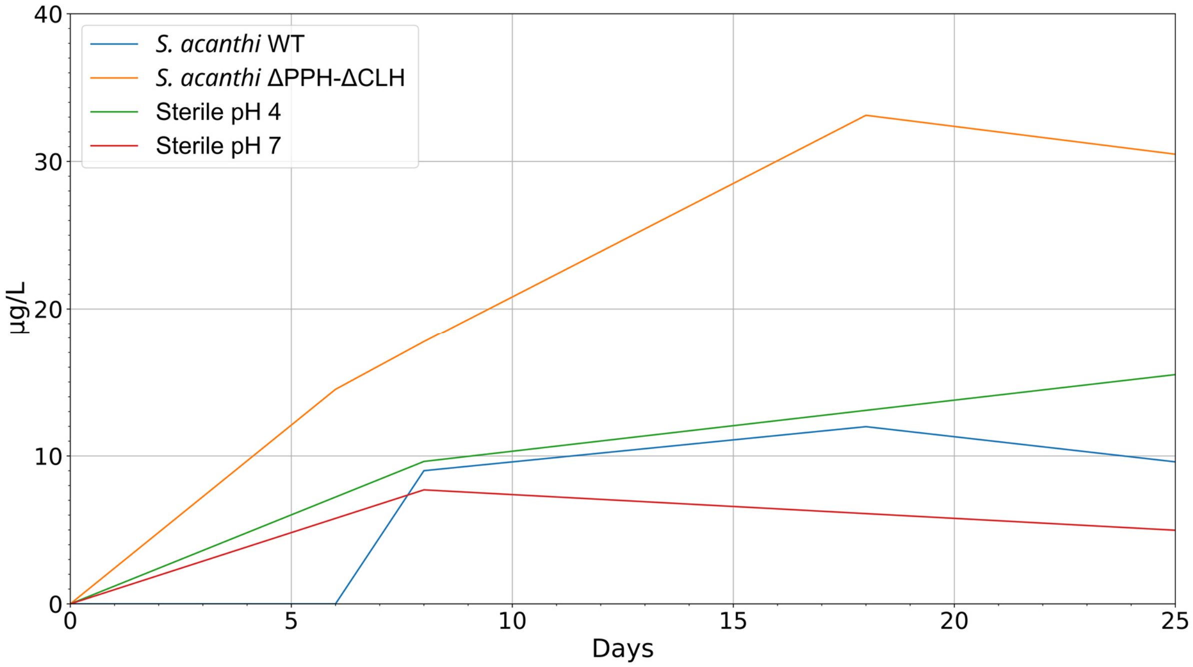


**Supplementary Fig. S6:** HPLC-based monitoring of Pheide *a*

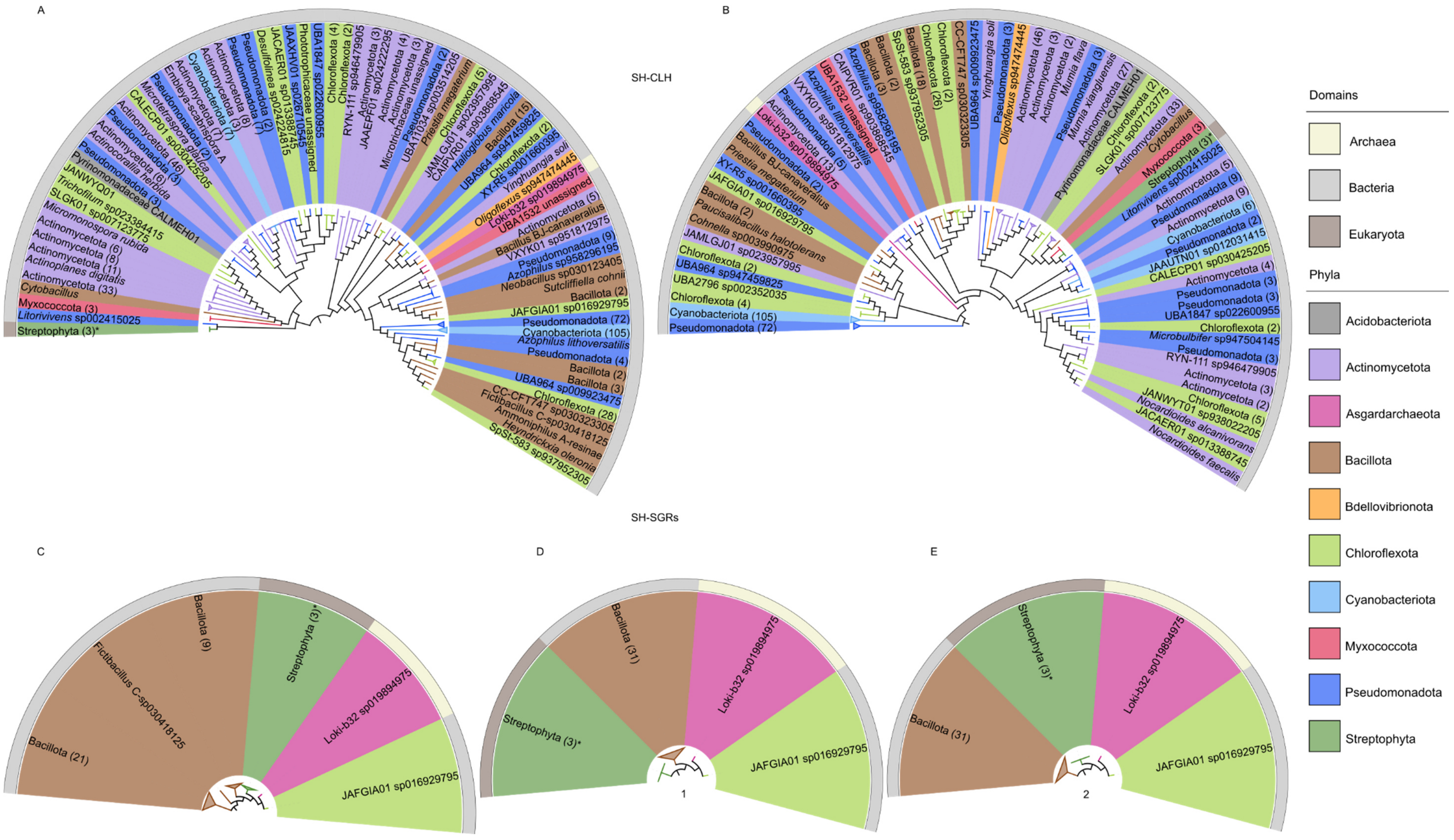
A
B
SH-CLH
Streptophyta (3)*
Litorivivens sp002415025
Myxococcota (3)
Cytobacillus
Actinomycetota (33)
Actinoplanes digitatis
Actinomycetota (11)
Actinomycetota (8)
Actinomycetota (6)
Micromonospora rubida
SLGK01 sp007123775
Trichofilum sp023384415
JANWYQ01
Pyrinomonadaceae CALMEH01
Pseudomonadota (3)
Actinocorallia herbida
Actinomycetota (6)
Pseudomonadota (3)
Actinomycetota (46)
CALECP01 sp030425205
Pseudomonadota (2)
Cyanobacteriota (7)
Actinomycetota (8)
Actinomycetota (3)
Pseudomonadota (2)
Pseudomonadota (2)
Desulfolinea sp022824815
JACAER01 sp013388745
JAAXHV01 sp026710545
Phototrophicaceae unassigned
UBA1847 sp022600955
Chloroflexota (4)
Chloroflexota (2)
RYN-111 sp946479905
JAAEPF01 sp024222295
Actinomycetota (3)
Actinomycetota (4)
Actinomycetota (3)
Microtrichaceae unassigned
UBA11034 sp003514205
Pseudomonadota (2)
Priestia megaterium
JAMLGJ01 sp023957995
CAIPVR01 sp903868545
Chloroflexota (5)
Haliangium maricola
Bacillota (15)
UBA964 sp947459825
XY-R5 sp001660395
Chloroflexota (2)
Yinghuangia soli
Oligoflexus sp947474445
Loki-b32 sp019894975
UBA1532 unassigned
Actinomycetota (5)
Bacillus BJ-canaveralius
VXYK01 sp951812975
Pseudomonadota (9)
Azophilus sp958296195
Neobacillus sp030123405
Sutcliffiella cohnii
Bacillota (2)
JAFGIA01 sp016929795
Pseudomonadota (72)
Cyanobacteriota (105)
Azophilus lithoversatilis
Pseudomonadota (4)
Bacillota (2)
Bacillota (3)
UBA964 sp009923475
Chloroflexota (28)
CC-CFT747 sp030323305
Fictibacillus C-sp030418125
Ammoniphilus A-resinae
Heyndrickxia oleronia
SpSt-583 sp937952305
Pseudomonadota (72)
Cyanobacteriota (105)
Chloroflexota (4)
UBA2796 sp002352035
UBA964 sp947459825
Chloroflexota (2)
JAMLGJ01 sp023957995
Cohnella sp003990975
Paucisalibacillus halotolerans
Bacillota (2)
JAFGIA01 sp016929795
XY-R5 sp001660395
Priestia megaterium
Bacillus BJ-canaveralius
Pseudomonadota (2)
Loki-b32 sp019894975
VXYK01 sp951812975
Actinomycetota (13)
Pseudomonadota (5)
Azophilus lithoversatilis
UBA1532 unassigned
CAIPVR01 sp903868545
Azophilus sp958296195
Bacillota (3)
Bacillota (2)
SpSt-583 sp937952305
Bacillota (18)
Chloroflexota (26)
Chloroflexota (2)
CC-CFT747 sp030323305
UBA964 sp009923475
Yinghuangia soli
Pseudomonadota (3)
Oligoflexus sp947474445
Actinomycetota (46)
Actinomycetota (3)
Actinomycetota (2)
Mumia flava
Pseudomonadota (3)
Mumia xiangluensis
Actinomycetota (27)
Pyrinomonadaceae CALMEH01
Chloroflexota (2)
SLGK01 sp007123775
Actinomycetota (33)
Cytobacillus
Myxococcota (3)
Streptophyta (3)*
Litorivivens sp002415025
Actinomycetota (5)
Pseudomonadota (9)
Actinomycetota (9)
Cyanobacteriota (6)
Pseudomonadota (2)
JAAUTN01 sp012031415
CALECP01 sp030425205
Actinomycetota (4)
Pseudomonadota (3)
Pseudomonadota (3)
UBA1847 sp022600955
Chloroflexota (2)
Microbulbifer sp947504145
Pseudomonadota (3)
RYN-111 sp946479905
Actinomycetota (3)
Actinomycetota (2)
Chloroflexota (5)
JANWYT01 sp938022205
Nocardioides alcanivorans
JACAER01 sp013388745
Nocardioides faecalis
SH-SGRs
C
Bacillota (21)
Fictibacillus C-sp030418125
Bacillota (9)
Streptophyta (3)*
Loki-b32 sp019894975
JAFGIA01 sp016929795
D
Streptophyta (3)*
Bacillota (31)
Loki-b32 sp019894975
JAFGIA01 sp016929795
1
E
Bacillota (31)
Streptophyta (3)*
Loki-b32 sp019894975
JAFGIA01 sp016929795
2
Domains
Archaea
Bacteria
Eukaryota
Phyla
Acidobacteriota
Actinomycetota
Asgardarchaeota
Bacillota
Bdellovibrionota
Chloroflexota
Cyanobacteriota
Myxococcota
Pseudomonadota
Streptophyta

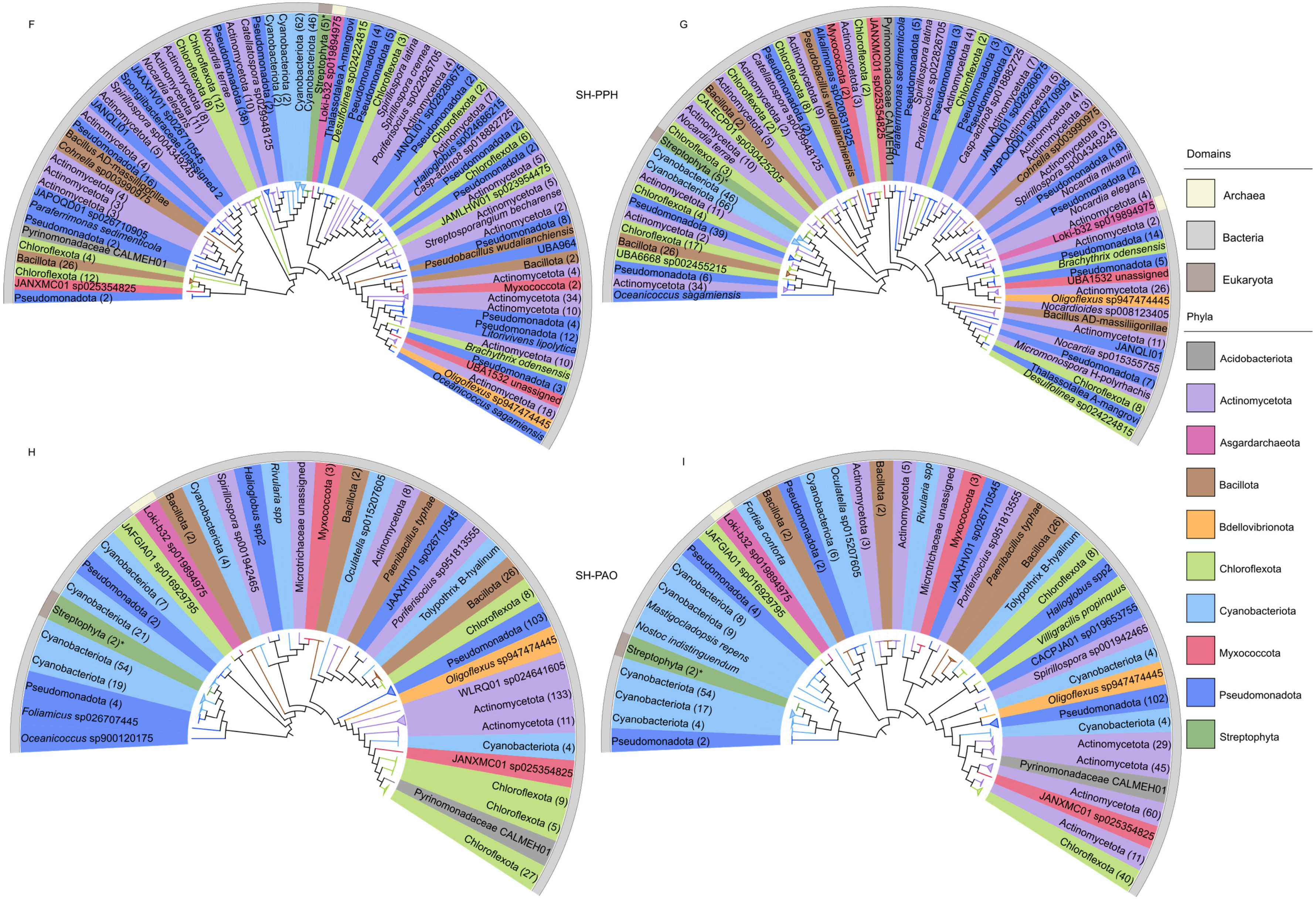
F
G
SH-PPH
H
I
SH-PAO
Domains
Archaea
Bacteria
Eukaryota
Phyla
Acidobacteriota
Actinomycetota
Asgardarchaeota
Bacillota
Bdellovibrionota
Chloroflexota
Cyanobacteriota
Myxococcota
Pseudomonadota
Streptophyta

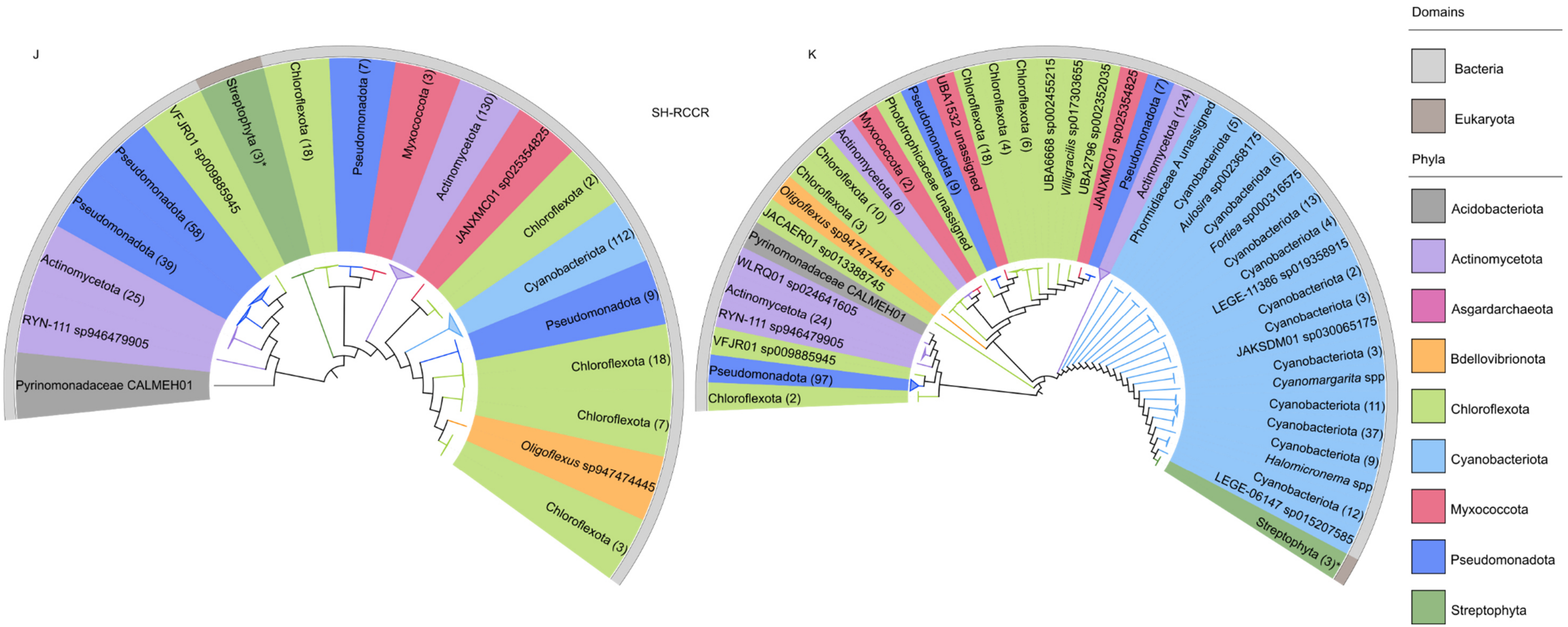


**Supplementary Fig. S7:** Maximum-likelihood phylogenies of predicted prokaryotic chlorophyll catabolic enzymes (SH-CCEs) in 469 taxa that contain complete SH-CCE sets. For each set of homologs, the top-scoring proteins were used to reconstruct the sequence (left) and the corresponding 3Di structure (right)-based phylogenies. Structures were predicted using AlphaFold3 [34] and the corresponding 3Di structural alignments were generated using FoldMason [35]. Multiple sequence alignment and refinement was done using The sequence and 3Di alignments were then trimmed with trimAl [37], and maximum-likelihood trees were inferred using IQ-TREE v3.0.1 [39] based on inferences for the best-fit models, including the 3Di substitution models [41] and an ultrafast bootstrap approximation with 1,000 replicates. **A**, **B**) SH-CLH maximum-likelihood trees comprise 469 prokaryotic and four plant homologs predicted under the models Q.PFAM+I+R7 (sequence) and 3DIPHY+F+R6 (3Di). **C**, **D**1,2) SH-SGRs maximum-likelihood trees, comprising thirty-three prokaryotic and six plant (Streptophyta) homologs, predicted under the models LG+G4 (sequence) and Q.3DI.AF+G4 (3Di: 1 and 2 differ by clock coefficient of variation(CVV of 0.7%)), **E**, **F**) SH-PPH maximum-likelihood trees include 469 prokaryotic and five plant (Streptophyta) homologs predicted under the models Q.PFAM+I+R6 (sequence) and Q.3DI.AF+F+I+R7 (3Di). **G**, **H**) SH-PAO maximum-likelihood trees consist of 469 prokaryotic and two plant (Streptophyta) homologs, predicted under the models LG+R7 (sequence) and Q.3DI.AF+F+R8 (3Di), and **I**, **J**) SH-RCCR maximum-likelihood trees contains 436 prokaryotic and three plant (Streptophyta) homologs, predicted under the models Q.PFAM+R8 (sequence) and Q.3DI.AF+F+R6 (3Di). An asterisk in plant branches indicates the number of plant species associated with the plant homologs. All trees were rooted using the minimal ancestor deviation (MAD) method [43]. In all phylogenies, we collapsed all monophyletic branches. We visualised the final trees in iTOL, with labels on the trees shaded according to the phyla of the corresponding taxa and provided corresponding colour keys in the legend.
